# Geometry-aware ligand-receptor analysis distinguishes interface association from spatial localization and reveals a continuum of tumor communication

**DOI:** 10.64898/2026.04.06.716708

**Authors:** Sally Yepes

## Abstract

Spatial transcriptomics enables inference of cell–cell communication through ligand–receptor (LR) interactions, but current prioritization strategies often rely on expression strength or interface-associated enrichment without explicitly modeling tissue geometry. As a result, interactions associated with population interfaces are frequently interpreted as spatially localized even when their underlying expression is broadly distributed.

Here, we present a geometry-aware framework for LR prioritization that explicitly separates interface structure from spatial localization within a locked and reproducible analysis pipeline. We quantify interface-associated communication using a distance-weighted boundary score defined on a spatial neighbor graph, evaluate interface specificity using a label-permutation null model that preserves spatial geometry, and compute an LR-specific localization score that captures the proximity of ligand and receptor expression to the corresponding interface. This framework distinguishes interface-associated compatibility from interaction-level spatial concentration.

Across spatial transcriptomics datasets from breast cancer, colorectal cancer, melanoma, and pancreatic ductal adenocarcinoma, interface-aware ranking consistently recovers pathway families associated with extracellular matrix, adhesion, inflammatory, and immune-related processes. However, interface enrichment frequently shows limited separation from the null model, indicating that interface structure alone does not establish spatial specificity. Incorporating geometric localization substantially alters LR prioritization, distinguishing interactions that are concentrated near interfaces from those that are more diffusely distributed.

Under a fixed, deterministic pipeline applied identically across datasets without parameter tuning, discrete spatial communication regimes were not reproducibly recovered. Instead, variation across samples is more consistently captured as continuous differences in geometry-aware attenuation, reflecting the degree to which inferred interactions are spatially constrained by tissue architecture.

Together, these results demonstrate that interface-associated enrichment and spatial localization are distinct properties of inferred LR interactions, and that accurate interpretation of spatial communication requires explicit modeling of tissue geometry. Under this framework, tumor communication is more consistently described as a continuum of spatial constraint.

## Introduction

Spatial transcriptomics enables the study of cellular organization and cell–cell communication within intact tissue architecture (Ståhl et al., 2016; Dries et al., 2021). A common approach to inferring intercellular communication relies on ligand–receptor (LR) interactions, where expression of a ligand in one cell population and its cognate receptor in another suggests potential signaling relationships. Methods such as CellPhoneDB and CellChat prioritize LR interactions using expression-derived measures and permutation-based testing, enabling systematic reconstruction of communication networks from transcriptomic data (Efremova et al., 2020; Jin et al., 2021).

Despite these advances, most LR prioritization approaches do not explicitly model spatial organization. As a result, interactions are often prioritized based on expression magnitude or interface-associated enrichment, even when the corresponding sender and receiver populations are spatially dispersed rather than concentrated near shared interfaces. This creates a fundamental interpretive problem: interface-associated signal does not imply spatial localization. Interactions may be compatible with structured adjacency between populations while remaining broadly distributed across the tissue.

Recent work has incorporated spatial context into transcriptomic analysis through spatially variable gene detection and neighborhood-based modeling (Svensson et al., 2018; Dries et al., 2021). However, explicit integration of tissue geometry into LR prioritization remains limited. In particular, existing approaches typically do not distinguish between population-level interface structure and interaction-level spatial deployment of gene expression. As a consequence, it remains unclear whether prioritized LR interactions reflect structured spatial organization or arise from properties of label arrangement and expression abundance.

A related question concerns how spatial communication varies across tumors. One possibility is that tumors can be categorized into discrete spatial communication regimes, such as localized or diffuse signaling. Alternatively, tumors may vary continuously in the degree to which inferred communication is constrained by tissue geometry. Resolving this requires a standardized analytical framework applied consistently across samples without dataset-specific parameter tuning, enabling separation of biological variation from analytical effects.

To address these challenges, we developed a geometry-aware framework for LR prioritization that explicitly separates interface structure from spatial localization under a locked and reproducible analysis pipeline, as illustrated in Figure 1. First, we quantify interface-associated communication using a distance-weighted boundary score defined on a spatial neighbor graph, capturing structured adjacency between annotated cell populations. Second, we evaluate interface specificity using a label-permutation null model that preserves spatial geometry while randomizing population labels, allowing assessment of whether observed interface structure exceeds randomized expectation. Third, we compute an LR-specific localization score that measures the proximity of ligand and receptor expression to the corresponding interface, distinguishing interface-associated from spatially concentrated signaling.

**Figure 1.**
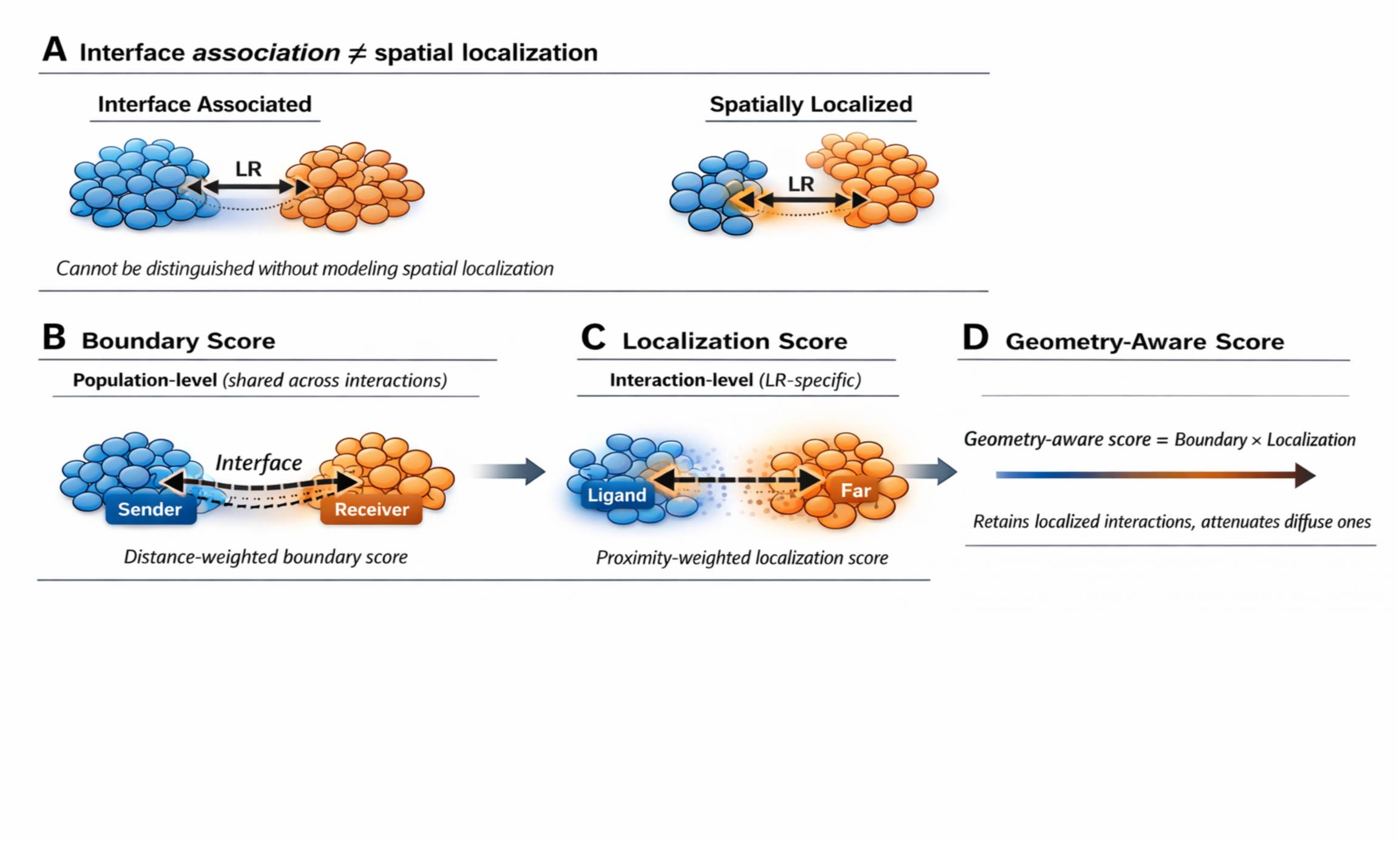
Geometry-aware framework separates interface structure from spatial localization. (A) Interface association ≠ spatial localization. Interactions associated with population interfaces may be either spatially diffuse or spatially localized, which cannot be distinguished without explicitly modeling spatial deployment. (B) Interface structure is quantified using a distance-weighted boundary score defined on cross-label edges in a spatial graph and represents a population-level property shared across interactions mapped to the same sender–receiver pair. (C) Spatial localization is quantified using an interaction-specific score based on the proximity of ligand and receptor expression to the corresponding interface, capturing the spatial deployment of signaling relative to tissue geometry. (D) The geometry-aware score integrates interface structure and spatial localization (Geometry-aware score = Boundary score × Localization score), retaining interactions that are spatially concentrated near interfaces and attenuating those that are more diffusely distributed.

**Figure 2.**
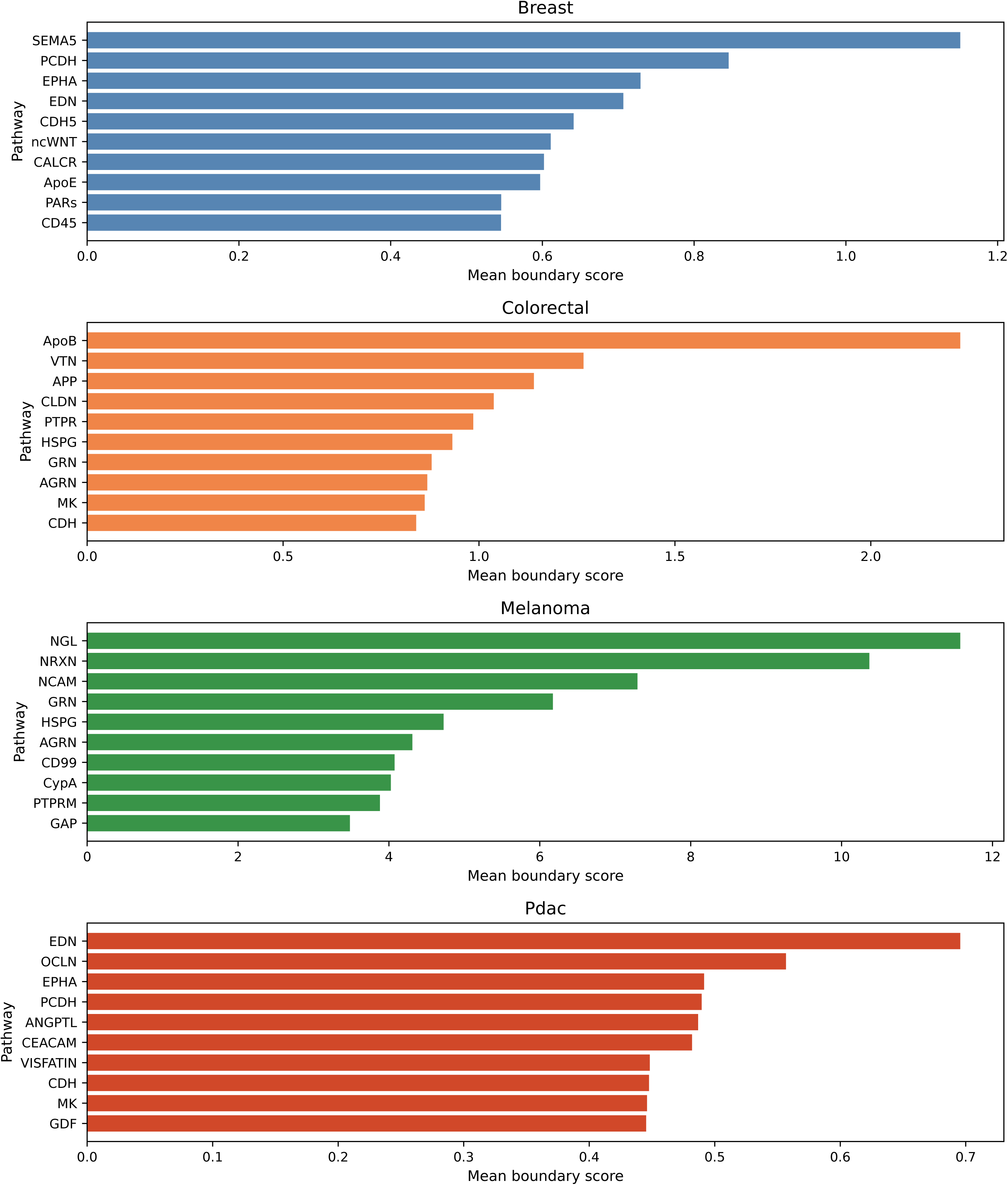
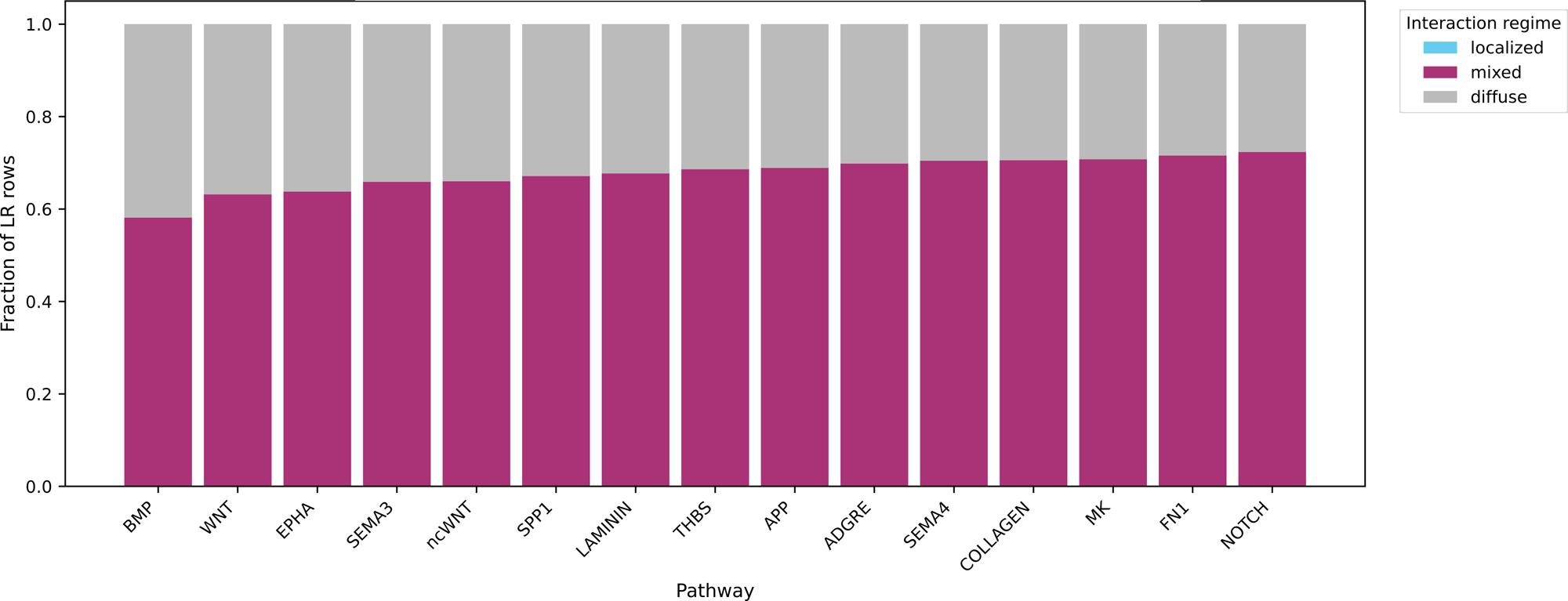
Interface-aware ranking recovers biologically plausible signaling programs across tumor tissues. A. Top pathways per tissue Top-ranked pathway families based on mean interface-associated boundary score within each tissue. Interface-aware ranking prioritizes pathway categories commonly associated with tumor–stroma and tumor–immune interactions, including extracellular matrix, adhesion, inflammatory, and immune-related processes. B. Regime composition of recurrent pathways Stacked fractions of interaction classifications (localized, mixed, diffuse) across recurrent pathway families. Most pathways are dominated by mixed classifications under the predefined thresholding scheme, with contributions from diffuse interactions and a limited fraction of strongly localized interactions, indicating that interface-associated enrichment does not necessarily correspond to spatial localization.

We applied this framework across spatial transcriptomics datasets from breast cancer, colorectal cancer, melanoma, and pancreatic ductal adenocarcinoma using an identical pipeline with fixed parameters and controlled stochastic components. Because the analytical framework is held constant across datasets, differences observed between samples reflect biological variation rather than dataset-specific tuning.

Using this approach, we show that interface-aware ranking recovers biologically plausible signaling programs but that interface enrichment alone frequently shows limited separation from randomized expectation. Incorporating geometric localization substantially alters LR prioritization, distinguishing interactions that are spatially concentrated near interfaces from those that are more diffusely distributed. Under this controlled framework, discrete spatial communication regimes are not reproducibly recovered. Instead, tumor samples vary along a continuum of spatial constraint, reflecting graded differences in the extent to which inferred interactions are localized relative to tissue architecture.

Together, these results establish that interface-associated enrichment and spatial localization are distinct properties of inferred LR interactions and demonstrate that accurate interpretation of spatial communication requires explicit modeling of tissue geometry.

## Results

### Interface-aware ranking recovers biologically plausible signaling programs across tumor tissues

We first asked whether interface-aware prioritization recovers signaling programs consistent with known features of tumor tissue organization. Across spatial transcriptomics datasets from breast cancer, colorectal cancer, melanoma, and pancreatic ductal adenocarcinoma, top-ranked ligand–receptor (LR) interactions recurrently involved extracellular matrix (ECM) signaling, adhesion pathways, inflammatory mediators, and immune-associated interactions. Representative examples included collagen- and laminin-associated interactions, FN1-related adhesion programs, MIF-associated signaling, and PPIA–BSG interactions, all of which have established roles in tumor–stroma and tumor–immune communication (Hynes, 2009; Hanahan and Weinberg, 2011; Binnewies et al., 2018).

Because the boundary score is defined on cross-label adjacency in a spatial neighbor graph, high-ranking interactions reflect structured interfaces between annotated populations rather than expression magnitude alone. Interface-aware ranking therefore captures aspects of tissue organization that are not accessible through abundance-based prioritization.

However, recovery of biologically plausible pathway families does not establish spatial localization. Many of these signaling programs are broadly expressed within contributing populations. Thus, interface-aware ranking identifies interactions compatible with tissue structure but does not determine whether they are spatially concentrated near the corresponding interface.

### Interface enrichment does not imply spatial specificity

We next evaluated whether strong interface-associated signal provides evidence of spatial specificity. Incorporating interface structure substantially altered LR prioritization relative to expression-based approaches, confirming that spatial adjacency influences which interactions are emphasized. However, interface-associated signal alone does not establish spatial specificity under randomized expectation.

To assess this, we compared observed boundary scores with a label-permutation null model that preserves the spatial graph and gene expression matrix while randomizing population labels. Across tissues, many interfaces with substantial boundary scores showed only limited separation from their null distributions (Figure 3A–B). Moreover, large boundary scores did not consistently correspond to strong deviation from the null model (Figure 3C). These results indicate that interface enrichment is common but often only weakly informative about spatial specificity.

**Figure 3.**
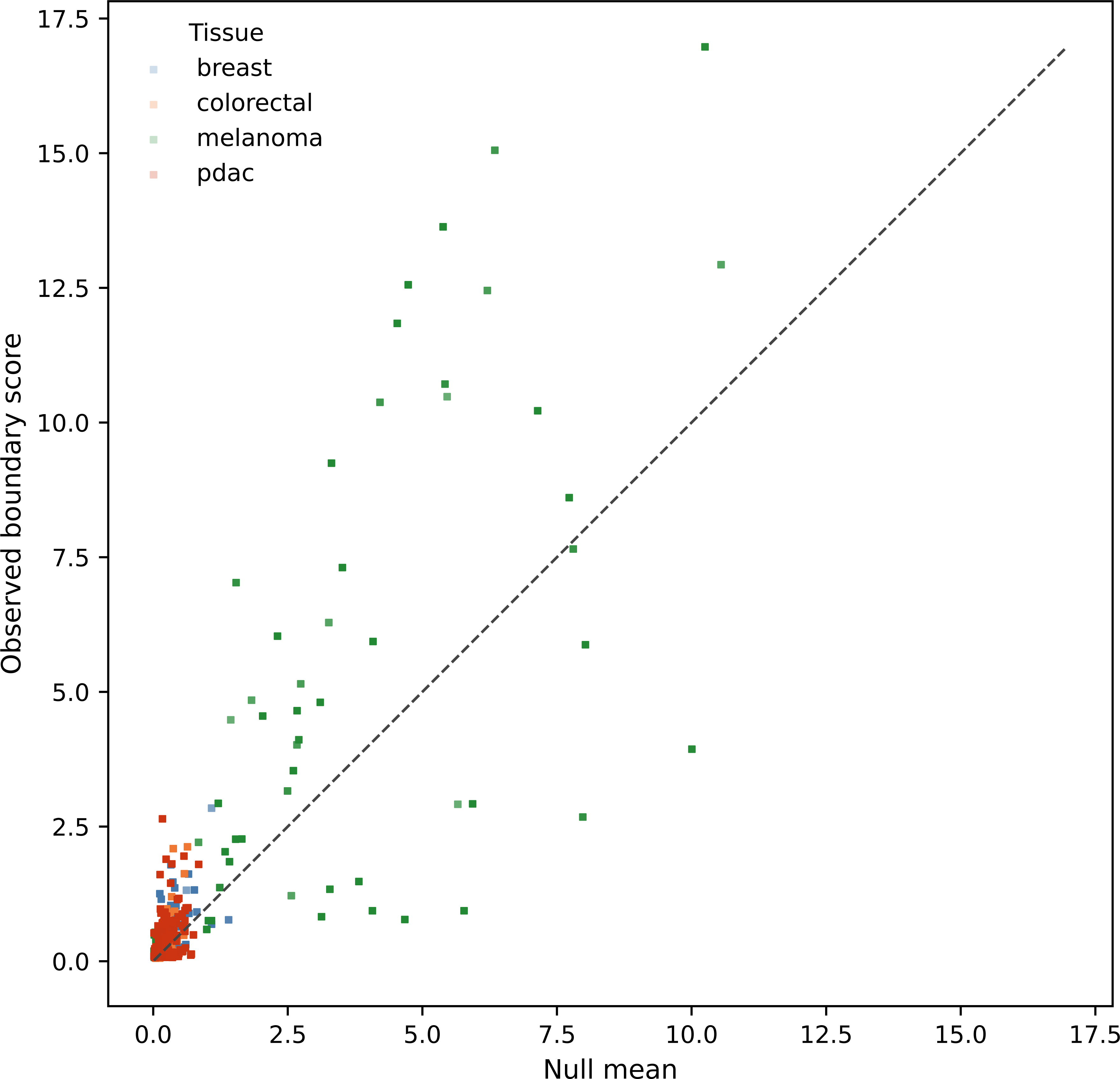
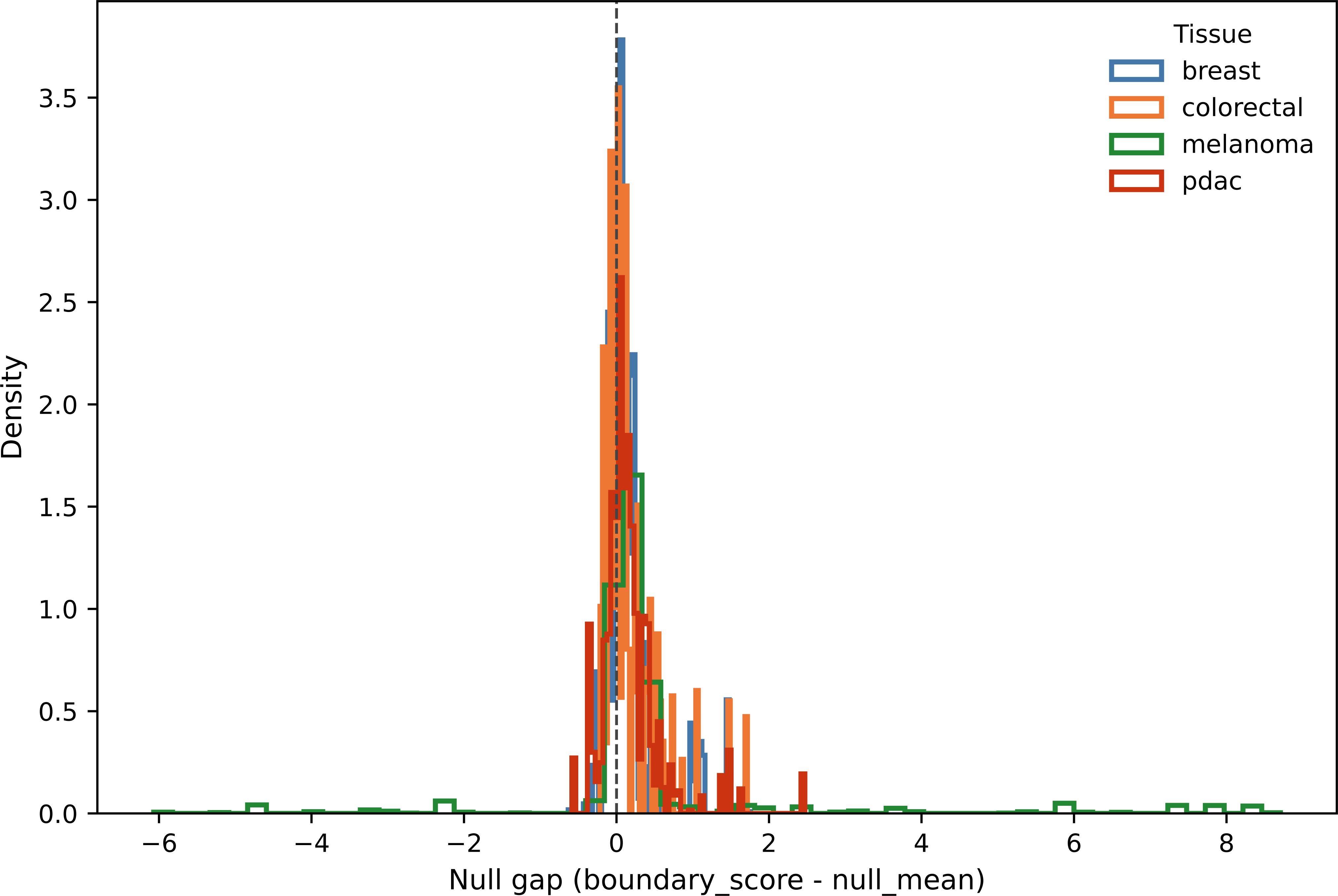
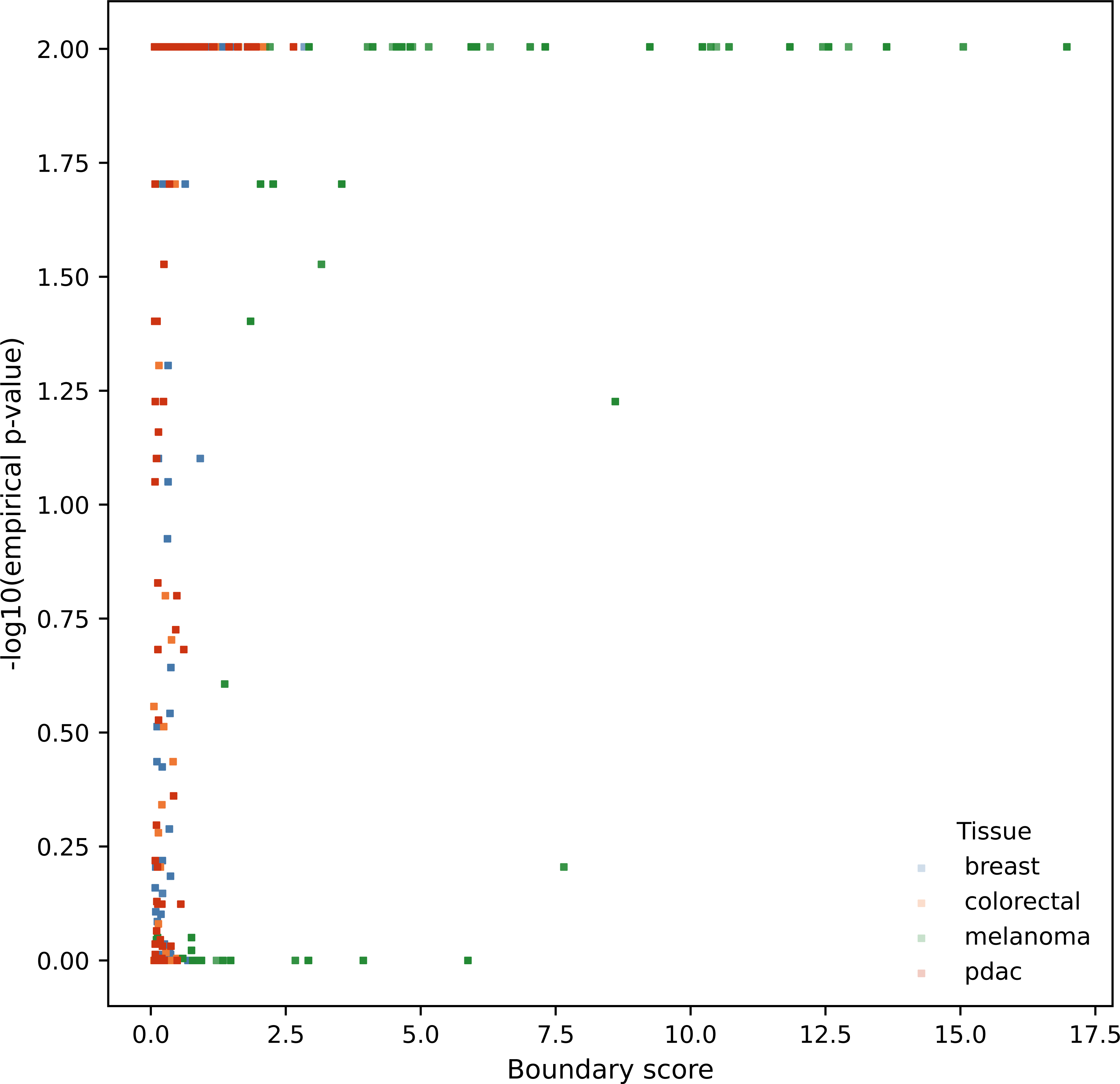
Interface enrichment does not imply spatial specificity. A. Observed versus null interface enrichment Comparison of observed boundary scores with mean null scores from label-permutation. Many interfaces with substantial boundary scores show only limited separation from randomized expectation, indicating that interface-associated enrichment alone does not establish spatial specificity. B. Distribution of null gaps Distributions of the difference between observed and null boundary scores across tissues. Values are broadly centered with substantial variability, reflecting heterogeneous but often modest deviation from randomized label organization. C. Effect size versus statistical significance Relationship between boundary score magnitude and empirical p-value derived from the label-permutation null model. Large boundary scores do not consistently correspond to strong deviation from null expectation, reinforcing that interface strength is not sufficient evidence of spatial specificity.

Taken together, these observations demonstrate that interface structure and spatial specificity are distinct properties. The boundary score captures structured adjacency between populations, whereas the null model evaluates whether that structure exceeds randomized label organization. Neither measure directly assesses whether a given LR interaction is spatially concentrated near the relevant interface.

### Geometry-aware refinement distinguishes interface-associated from interface-localized signaling

To distinguish interface-associated compatibility from spatially localized signaling, we applied an LR-specific geometry-aware refinement that evaluates the proximity of ligand and receptor expression to the corresponding interface, as conceptually illustrated in Figure 1.

Incorporating this refinement substantially altered LR prioritization across datasets. Interactions whose expression was concentrated near the relevant interface retained relatively high geometry-aware scores, whereas interactions with broader spatial deployment were attenuated (Figure 4A). Importantly, these differences emerged among interactions mapped to the same interface: LR pairs sharing an identical boundary score exhibited distinct geometry-aware retention depending on how expression was distributed relative to the interface.

**Figure 4.**
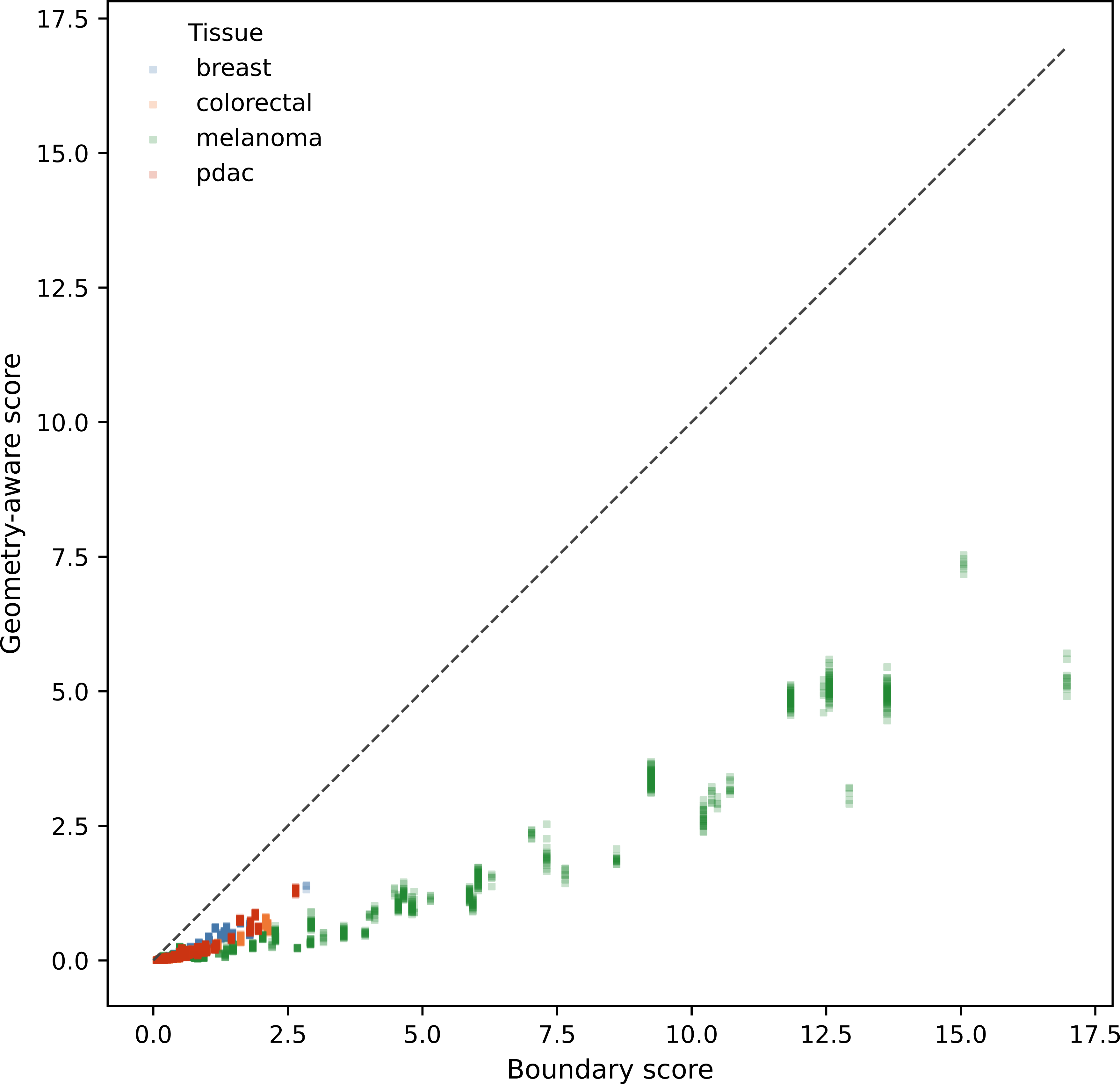
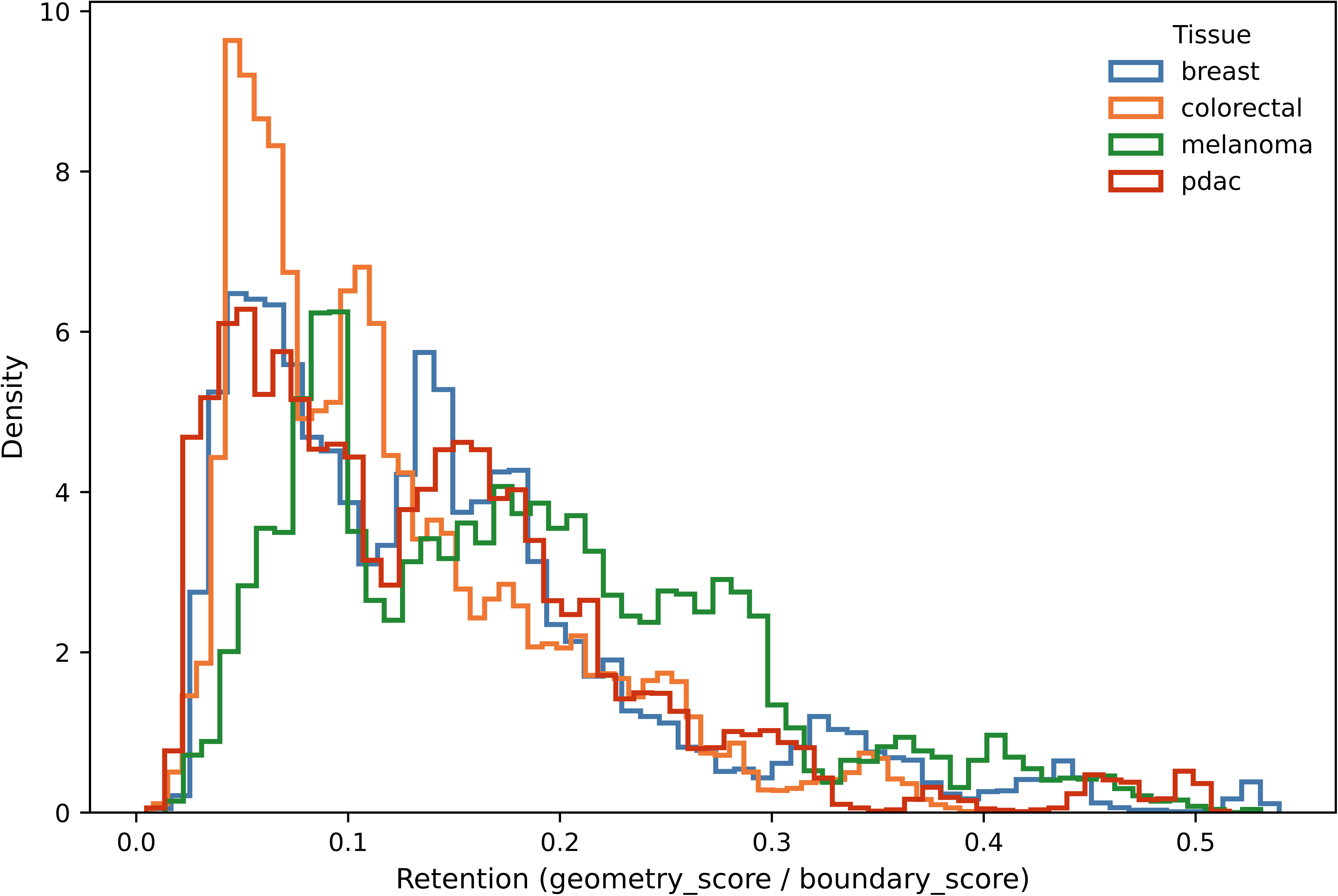
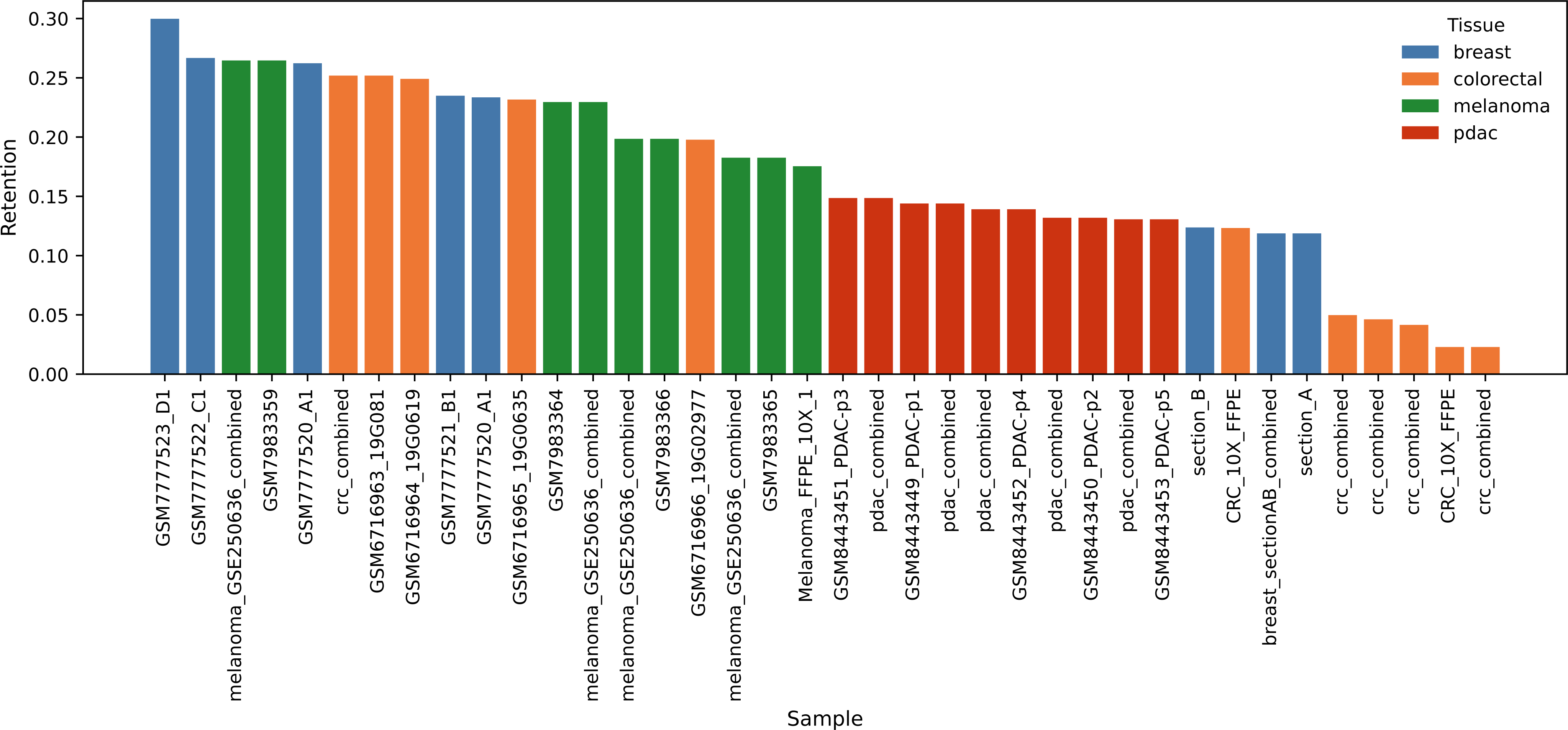
Geometry-aware refinement distinguishes interface-associated from spatially localized signaling. A. Boundary versus geometry-aware scores Comparison of interface-level boundary scores with geometry-aware scores across LR interactions. Geometry-aware scores are generally attenuated relative to boundary scores, reflecting penalization of interactions with diffuse spatial expression. B. Retention distribution Distribution of retention values (geometry-aware score divided by boundary score) across interactions. Most interactions exhibit partial retention, indicating that only a subset of interface-associated interactions remain spatially concentrated after accounting for geometric localization. C. Sample-level retention Retention summarized at the sample level. Samples exhibit continuous variation in retention without clear separation into discrete groups, consistent with graded differences in spatial constraint.

This separation clarifies the roles of the two components of the framework. The boundary score is a population-level measure shared across interactions assigned to the same sender–receiver pair, whereas localization operates at the level of individual LR interactions and penalizes diffuse expression. The geometry-aware score therefore integrates interface structure with spatial deployment and is best interpreted as a prioritization metric rather than a direct estimate of signaling strength.

Across datasets, attenuation after geometry-aware refinement was common (Figure 4B). Most interactions exhibited partial retention, indicating that only a subset of interface-associated interactions remain strongly localized after accounting for geometric proximity. This demonstrates that interface-associated signaling frequently reflects compatibility with tissue structure without corresponding spatial concentration.

### Discrete spatial communication regimes are not reproducibly recovered under a fixed framework

We next asked whether tumors can be organized into discrete spatial communication regimes based on the relationship between interface-associated and geometry-aware signaling. To do this, we summarized each interaction using fixed thresholds applied to retention and null-related metrics and classified interactions as localized, diffuse, or mixed. These classifications were aggregated into sample-level summaries under an identical pipeline across datasets.

Under this predefined and fixed thresholding scheme, discrete regime classes were not reproducibly recovered. Sample-level summaries were dominated by mixed classifications, and strongly localized interactions represented only a small fraction of the total across datasets (Figure 2B, Figure 5). Differences between samples were therefore expressed primarily as quantitative shifts in retention and diffuse fractions rather than as stable separation into distinct categories.

**Figure 5.**
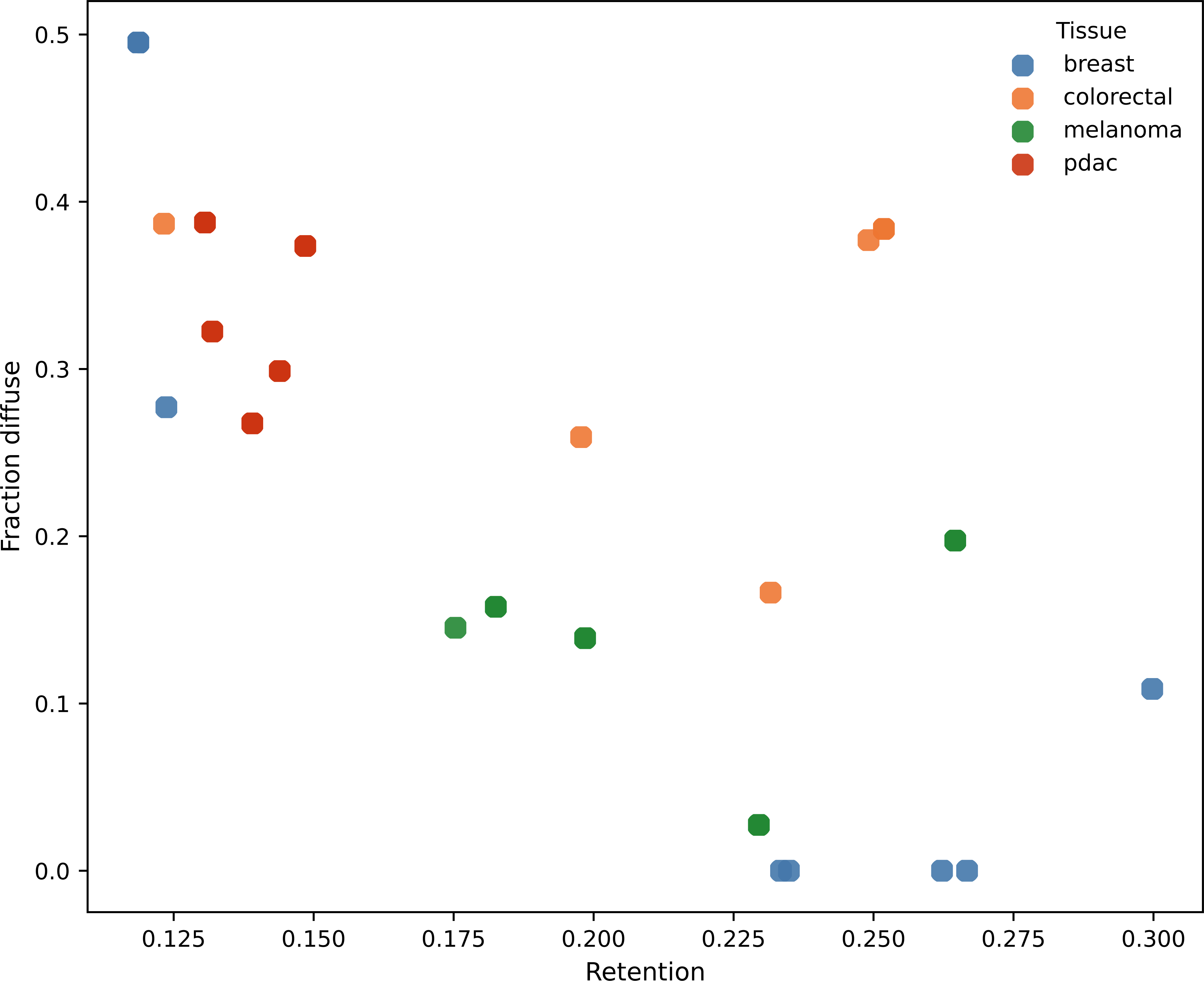
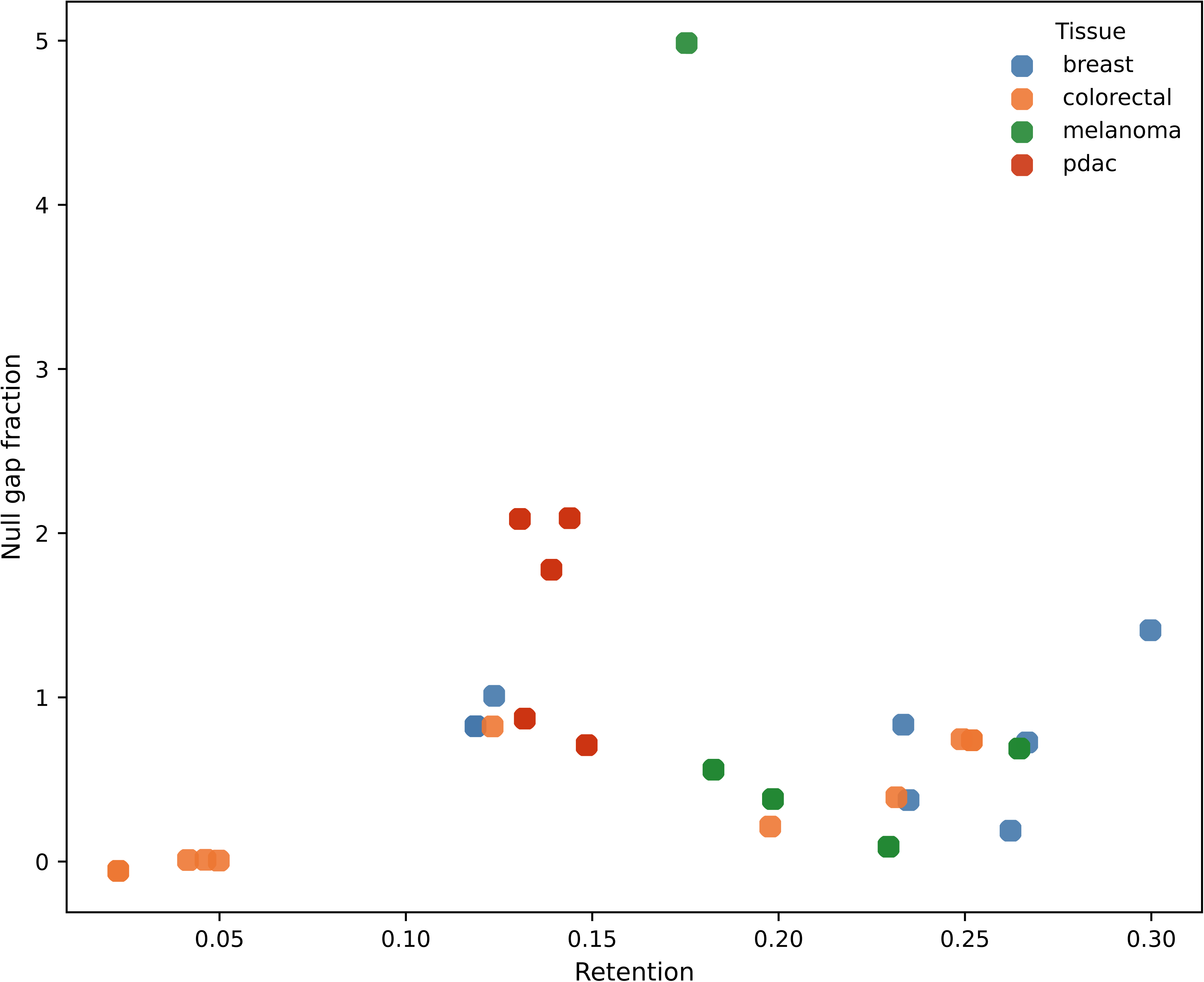
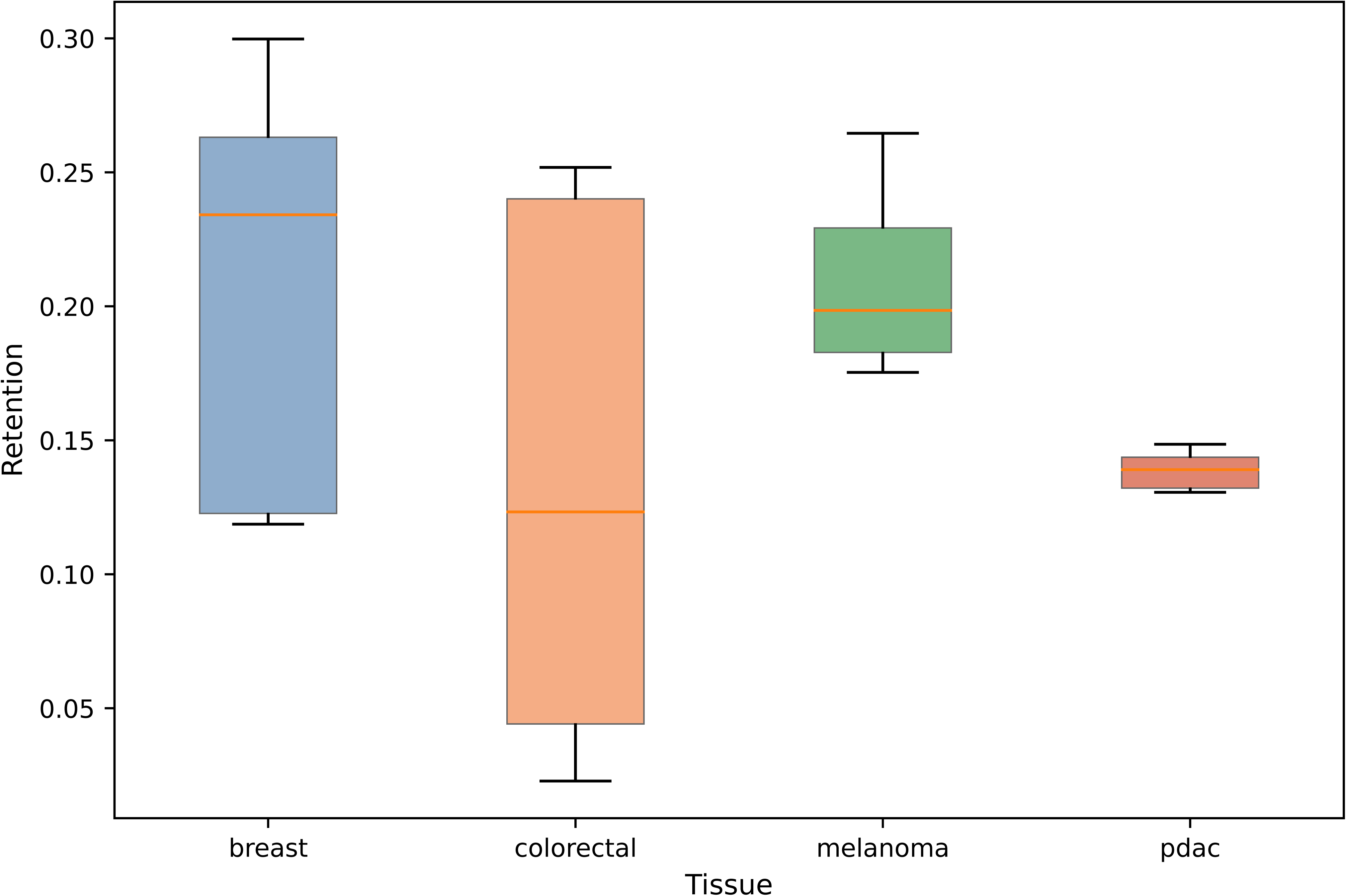
Tumor samples vary along a continuum of spatial constraint. A. Retention versus diffuse fraction Sample-level relationship between mean retention and fraction of interactions classified as diffuse. Samples occupy a continuous range without clear clustering into discrete regimes. B. Retention versus null-related metric Alternative projection showing retention in relation to null-derived quantities. The same continuous structure is observed, reinforcing the absence of stable threshold-defined classes. C. Retention distributions by tissue Distribution of sample-level retention across tissues, highlighting overlapping ranges and continuous variation rather than discrete separation.

Because classification is performed under fixed thresholds without dataset-specific tuning, this analysis evaluates the stability of a predefined regime formalization rather than optimizing for separation. Within this controlled framework, the absence of stable classes indicates that discrete spatial communication regimes are not strongly supported by the data.

### Tumor samples vary along a continuum of spatial constraint

Although discrete regimes were not recovered, a consistent pattern emerged across the cohort: samples varied continuously in the degree to which geometry-aware refinement attenuated interface-associated interactions.

At one end of this spectrum, samples retained a larger fraction of their interface-associated signal after geometry-aware refinement, indicating stronger alignment between population interfaces and the spatial deployment of ligand and receptor expression. At the other end, samples exhibited greater attenuation and higher diffuse fractions, consistent with interface-associated signaling that is more broadly distributed across contributing populations. Rather than forming discrete groups, samples occupied overlapping positions along this axis of spatial constraint (Figure 5A–C).

Because all datasets were processed under an identical pipeline with fixed parameters and controlled stochastic components, this continuous variation is unlikely to arise from analytical flexibility or parameter tuning. Instead, it reflects intrinsic differences in how spatial communication is organized across tissues.

### Conserved pathway families are deployed under different degrees of spatial constraint

Finally, we examined whether recurrent signaling pathways are preserved across tissues despite differences in spatial organization. Across the cohort, several pathway families consistently appeared among top-ranked interactions, including ECM-associated pathways, adhesion programs, inflammatory mediators, and immune-related interactions (Hynes, 2009; Hanahan and Weinberg, 2011; Binnewies et al., 2018).

However, the extent to which these pathways remained prioritized after geometry-aware refinement differed across samples and tissues (Figure 6). In some contexts, pathway-associated interactions retained a substantial fraction of their interface-associated signal, consistent with expression concentrated near interfaces. In others, the same pathway families exhibited stronger attenuation, indicating broader spatial deployment.

**Figure 6.**
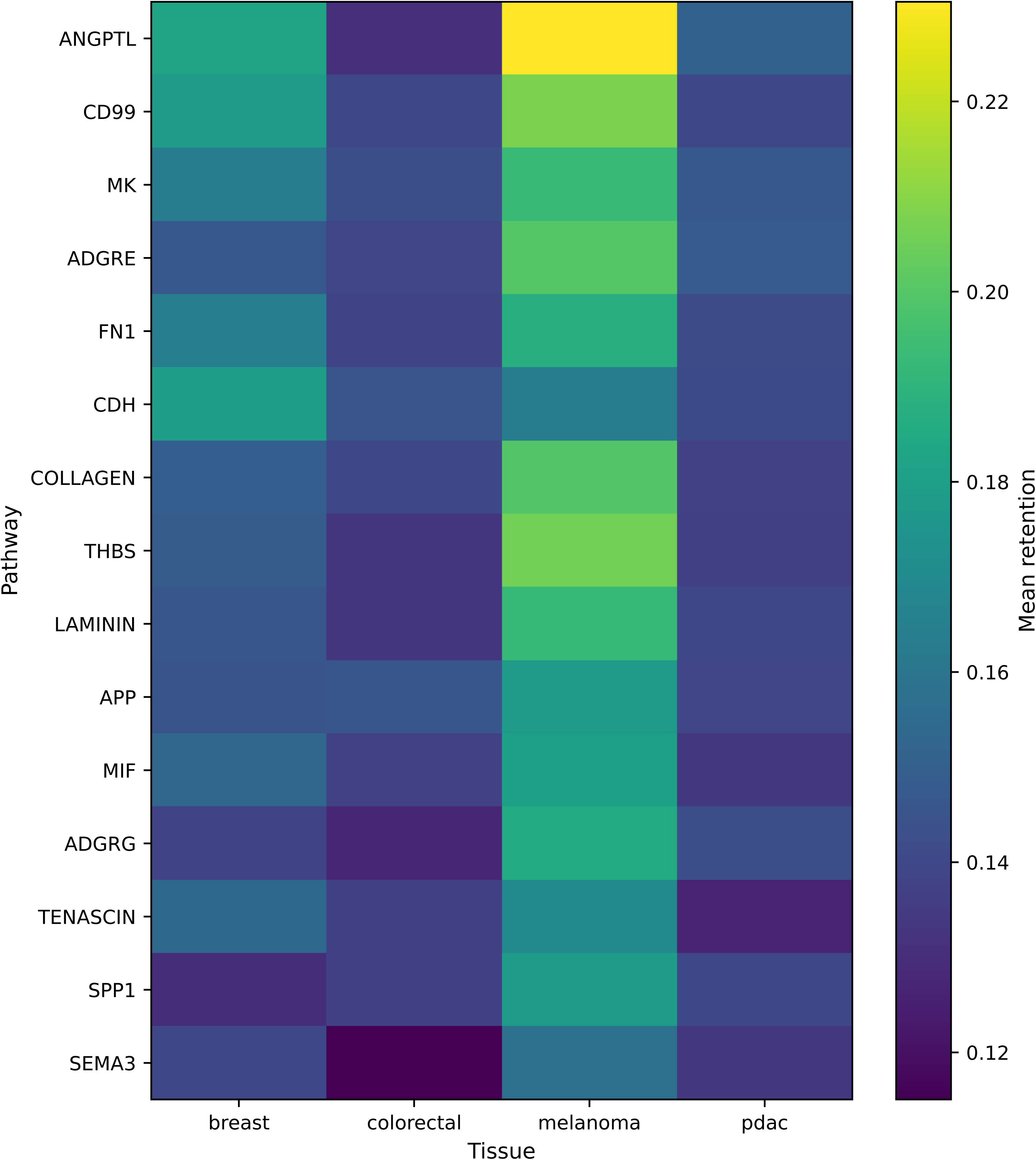
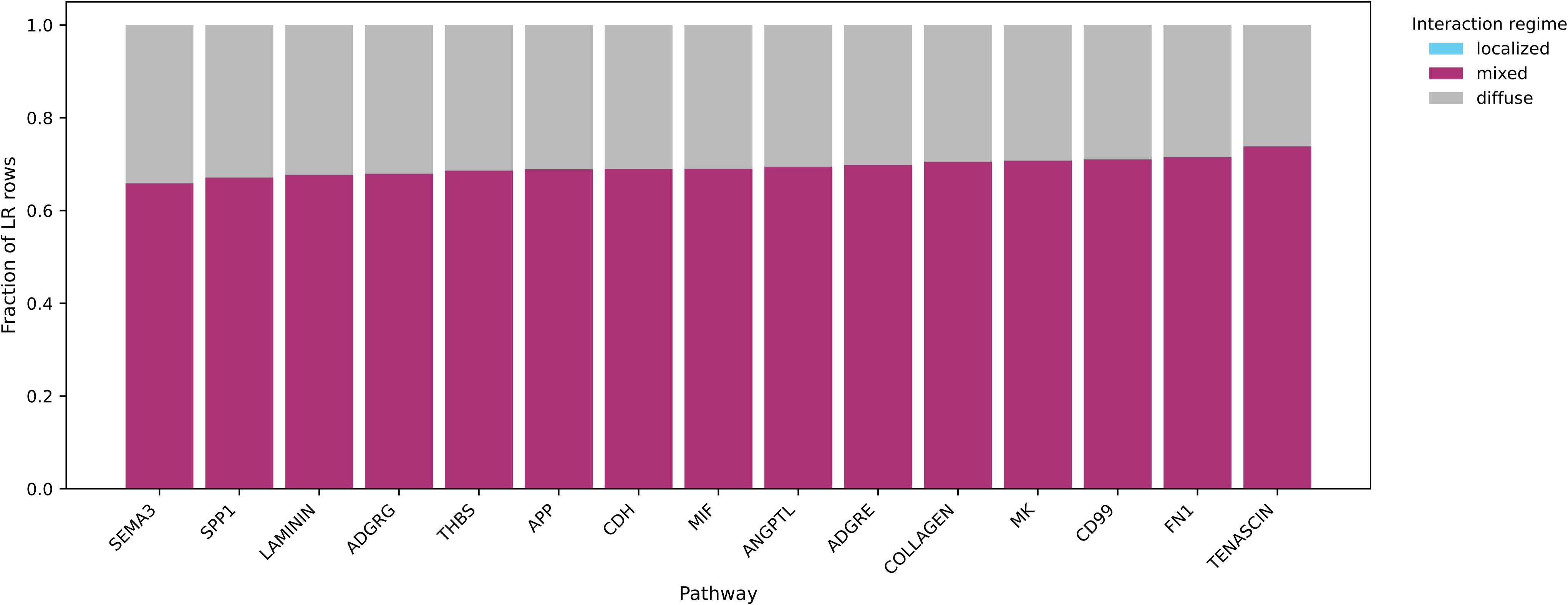
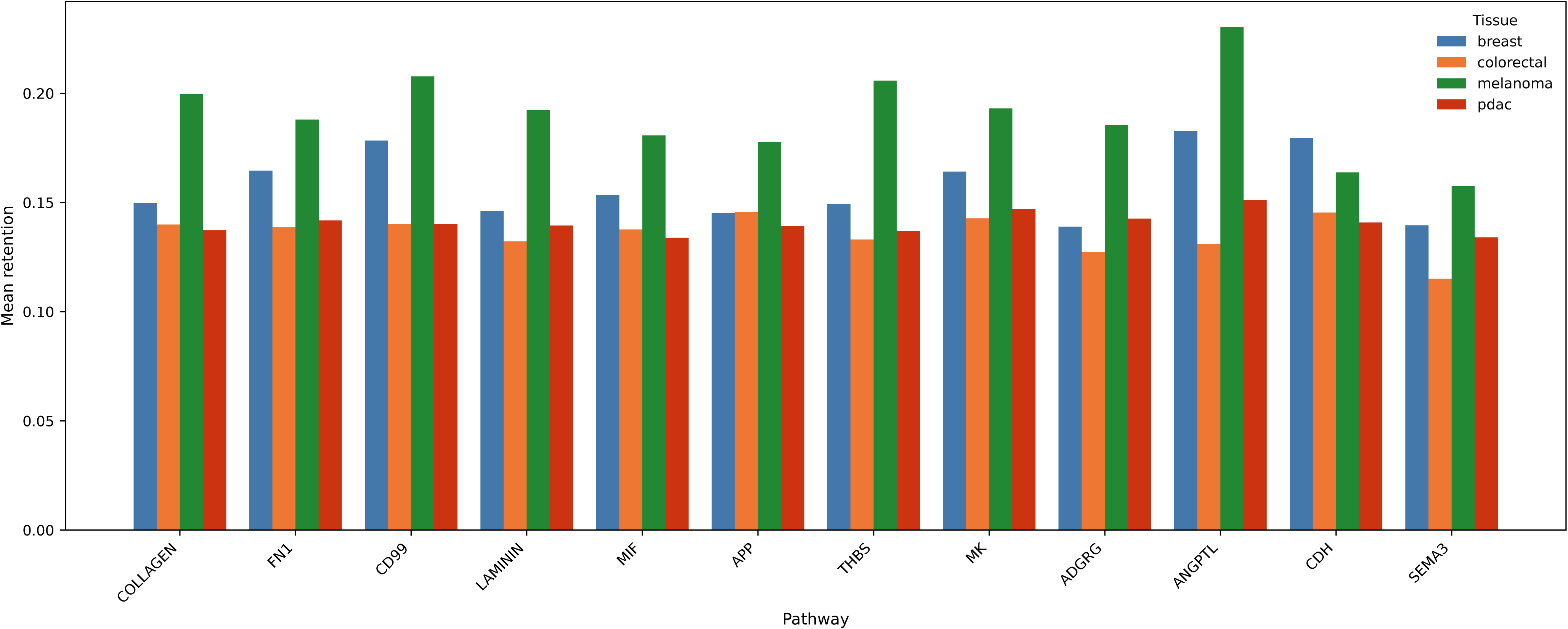
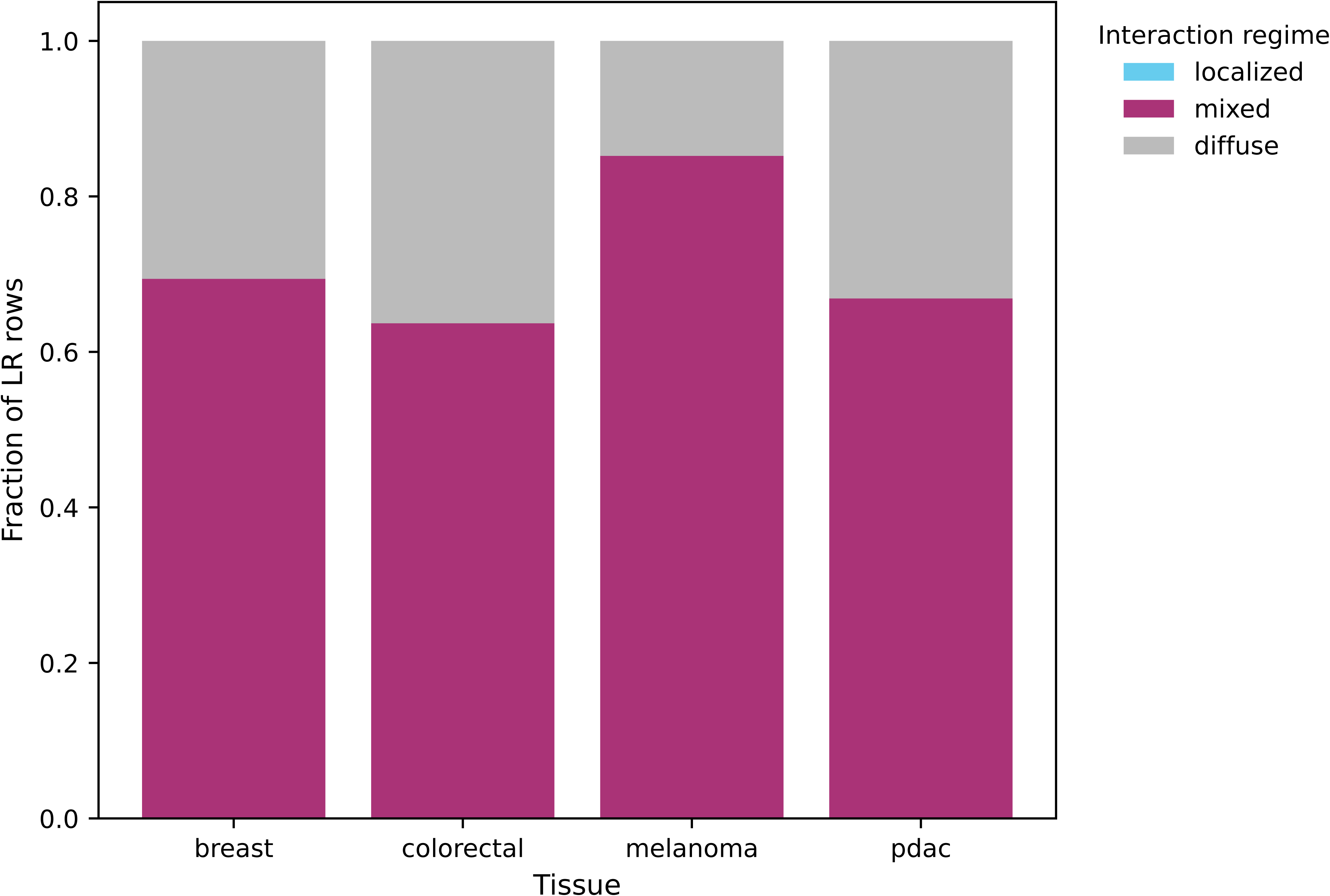
Conserved pathway families are deployed under different degrees of spatial constraint. A. Pathway retention across tissues Heatmap of mean retention for recurrent pathway families across tissues. Pathways are shared across tumor types but differ in the extent to which their associated interactions remain spatially concentrated near interfaces. B. Regime composition of conserved pathways Distribution of localized, mixed, and diffuse interaction classifications for conserved pathways across tissues. Differences reflect variation in spatial deployment rather than pathway identity. C. Tissue-level comparison of pathway retention Grouped comparison of retention for selected pathway families across tissues, highlighting variation in spatial constraint for shared signaling programs. D. Global regime composition by tissue Overall distribution of interaction classifications within each tissue. Mixed interactions dominate across tissues, with contributions from diffuse interactions and a limited fraction of strongly localized signal, consistent with continuous rather than discrete spatial organization.

These results indicate that conserved signaling programs are deployed under different geometric constraints across tumor tissues. Thus, cross-sample variation arises not only from pathway composition but also from how shared signaling programs are organized in space. Spatial constraint therefore represents an additional dimension of tumor organization beyond pathway identity alone.

Taken together, these analyses demonstrate that interface-aware prioritization recovers biologically meaningful signaling programs, but that their spatial interpretation depends critically on whether expression is concentrated near interfaces. Incorporating geometric localization refines LR prioritization by distinguishing interface-associated compatibility from interaction-level spatial concentration and reveals continuous variation in spatial constraint across tumor tissues.

## Discussion

A central conclusion of this study is that interface-associated enrichment and spatial localization are distinct properties of inferred ligand–receptor (LR) interactions and should not be treated as interchangeable. The interface-aware component of our framework consistently prioritizes interactions mapped to structured boundaries between annotated cell populations, recovering pathway families that are consistent with known tumor–stroma and tumor–immune biology (Hynes, 2009; Hanahan and Weinberg, 2011; Binnewies et al., 2018). However, strong interface-associated signal does not necessarily imply spatial specificity under a label-permutation null model, nor does it ensure that ligand and receptor expression are concentrated near the corresponding interface. Interpreting interface enrichment as evidence of localized signaling may therefore lead to systematic overinterpretation of spatial communication.

The geometry-aware refinement step addresses this limitation by explicitly modeling the spatial deployment of gene expression relative to interface structure. By separating population-level interface compatibility from interaction-level spatial concentration, as illustrated in Figure 1, the framework distinguishes interactions that are merely compatible with tissue architecture from those that are spatially localized. Because the boundary score is defined at the level of population pairs whereas localization is computed for individual LR interactions, the geometry-aware score integrates these components into a prioritization metric that reflects both structure and deployment without conflating them.

The cross-sample analysis further clarifies how spatial communication should be interpreted across tumor tissues. Under a fixed and deterministic analytical pipeline applied identically across datasets, discrete spatial communication regimes were not reproducibly recovered. Instead, variation between samples was expressed as continuous differences in geometry-aware attenuation and diffuse fractions. Because all datasets were processed under identical parameters without dataset-specific tuning, this continuous structure cannot be attributed to analytical flexibility or threshold optimization. Rather, it reflects intrinsic variation in the degree to which inferred communication is spatially constrained by tissue architecture.

This continuum-based interpretation provides a parsimonious alternative to regime-based descriptions of spatial communication. Rather than assigning tumors to discrete categories, the data support a quantitative view in which samples differ in the extent to which inferred interactions remain localized relative to population interfaces. This perspective is consistent with broader observations that biological organization often exhibits graded structure rather than sharply defined classes (Trapnell, 2015).

The observation that conserved pathway families are deployed under different degrees of spatial constraint further supports this interpretation. Extracellular matrix, adhesion, inflammatory, and immune-related pathways were consistently identified across tumor types, but differed in the extent to which their associated interactions remained spatially concentrated near interfaces (Hynes, 2009; Hanahan and Weinberg, 2011; Binnewies et al., 2018). This indicates that cross-tissue variation arises not only from which signaling pathways are active, but also from how those pathways are organized in space. Spatial constraint therefore represents an additional dimension of tumor organization that complements established axes such as cell-type composition and transcriptional state.

These findings have several practical implications for the analysis of spatial transcriptomics data. First, expression-based LR prioritization alone is insufficient to identify spatially plausible interactions. Second, interface-aware scoring captures structured adjacency between populations but should be interpreted as a property of tissue organization rather than as evidence of localized signaling. Third, label-permutation null models provide a principled way to evaluate whether interface structure exceeds randomized expectation, but operate at the population level and do not assess interaction-specific localization. Fourth, incorporating geometric proximity improves interpretability by distinguishing interactions that are spatially concentrated near interfaces from those that are more diffusely distributed. Together, these considerations support the use of multi-component frameworks that explicitly integrate expression, interface structure, and spatial deployment.

Several limitations should be considered. The null model evaluates interface structure at the level of population pairs and does not provide LR-specific significance estimates. The geometry-aware score is a relative prioritization metric and is not normalized across interfaces or tissues. Regime classification depends on predefined thresholds applied to retention and null-related metrics and therefore reflects a specific operational definition rather than an exhaustive exploration of possible discrete structures. In addition, the analysis depends on the quality and consistency of population labels and on how multimeric ligands and receptors are represented in expression data. Finally, the study is computational and does not include orthogonal experimental validation of prioritized interactions.

In summary, this study demonstrates that interface-associated enrichment alone can misrepresent the spatial organization of inferred LR interactions and that explicit modeling of tissue geometry is required to distinguish interface-associated compatibility from spatial localization. Under a controlled and reproducible analytical framework, spatial communication across tumor tissues is more consistently described as a continuum of spatial constraint rather than as discrete regimes. These results establish geometry-aware modeling as a necessary component of interpretable cell–cell communication analysis in spatial transcriptomics and position spatial constraint as a quantitative dimension of tumor organization.

## Methods

### Overview of the geometry-aware analysis framework

All analyses were performed using a locked and reproducible computational pipeline applied identically across datasets without dataset-specific parameter tuning, threshold adjustment, or model reconfiguration. The workflow consists of preprocessing, spatial graph construction, interface scoring, ligand–receptor (LR) mapping, null model evaluation, geometry-aware prioritization, and sample-level summarization. Each dataset is processed independently using the same analytical steps, and all stochastic components are controlled via fixed random seeds to ensure deterministic behavior.

To support reproducibility and modularity, each dataset is analyzed within a dedicated Jupyter notebook implementing the identical pipeline, while a separate workflow aggregates outputs to generate cohort-level figures. This structure ensures that all cross-sample differences arise from the data rather than analytical variation.

### Spatial transcriptomics preprocessing

Raw spatial transcriptomics count matrices were processed using Scanpy (Wolf et al., 2018). Counts were normalized by total counts per spot and log-transformed. Highly variable genes (n = 3,000) were selected using the Seurat flavor method (Hao et al., 2021), and principal component analysis (PCA) was performed using 30 components.

A k-nearest-neighbor (kNN) graph in expression space was constructed using 15 neighbors, and Leiden clustering was performed at resolution 0.6 (Traag et al., 2019). These steps provide a standardized low-dimensional representation and clustering backbone but are not directly used for LR scoring.

### Cell population labels for LR analysis

Ligand–receptor analysis was performed using CellChat-derived population labels (Jin et al., 2021). Cell population annotations were mapped to spatial barcodes to ensure alignment between spatial transcriptomics data and LR interaction tables. Only labels present in both the spatial dataset and the LR interaction table were retained.

These labels define the sender–receiver population pairs used for interface construction and LR mapping.

### Spatial graph construction

For each sample, a spatial k-nearest-neighbor graph was constructed from spot coordinates using Euclidean distance, with a fixed neighborhood size of six neighbors per spot. Edge weights correspond to Euclidean distances and define local spatial adjacency.

This graph provides the geometric substrate for both interface scoring and localization analysis.

### Interface definition and boundary score

Interfaces were defined as sets of edges in the spatial graph connecting spots assigned to different population labels. For each ordered pair of populations, all cross-label edges were collected.

Each edge was weighted by the inverse of its Euclidean distance, assigning greater contribution to closer spatial interactions. The boundary score for a population pair was defined as the sum of inverse-distance weights over all edges connecting the two populations. Interfaces supported by fewer than 20 cross-label edges were excluded to reduce instability from sparse boundaries.

The boundary score is an interface-level quantity shared across all LR interactions mapped to the corresponding population pair and reflects structured adjacency between populations rather than interaction-specific localization.

### Ligand–receptor interaction mapping

CellChat-derived LR interactions were filtered to retain entries with valid sender and receiver labels present in the spatial dataset. Each LR interaction was mapped to its corresponding ordered population pair, and the associated boundary score was assigned to all interactions mapped to that interface.

For multimeric ligands or receptors, expression was approximated using the minimum expression across detected subunits, providing a conservative estimate of complex activity when all components are not jointly observed.

### Label-permutation null model

To evaluate whether observed interface structure exceeds randomized expectation, a label-permutation null model was implemented. The spatial graph, edge distances, and gene expression matrix were held fixed, while population labels were randomly permuted across spatial locations, preserving label frequencies.

For each permutation, boundary scores were recomputed using the same distance-weighted formulation. This generates a null distribution for each population pair under randomized label organization within the fixed tissue geometry. Empirical p-values were computed by comparing observed boundary scores to the corresponding null distribution using 100 permutations.

This null model evaluates interface structure at the population level and does not provide LR-specific measures of spatial localization.

### Geometry-aware localization

To distinguish interface-associated compatibility from spatial localization, an LR-specific localization score was computed for each interaction.

For each interface, boundary-edge midpoints were used to represent interface geometry. For a given LR interaction, ligand expression was evaluated in sender spots and receptor expression in receiver spots. Distances from expressing spots to the nearest interface representation were computed and transformed into proximity weights using an exponential decay function.

The decay scale was defined for each interface as the median distance of its boundary edges, allowing localization to adapt to local geometric scale. Sender-side and receiver-side proximity scores were averaged to obtain a single LR-specific localization score.

### Geometry-aware prioritization

The geometry-aware score was defined as the product of the boundary score and the localization score:

### Geometry-aware score = Boundary score × Localization score

This formulation retains high values for interactions supported by strong interfaces and spatially concentrated expression, while attenuating interactions that are diffuse relative to the interface. Because the boundary score is interface-specific, cross-sample comparisons are performed using retention rather than raw geometry-aware values.

Retention was defined as the ratio of geometry-aware score to boundary score and represents the fraction of interface-associated signal preserved after accounting for geometric localization.

### Sample-level summarization and regime classification

Each LR interaction was summarized using retention and null-related metrics, including the difference between observed and mean null boundary scores. Interactions were classified as localized, diffuse, or mixed using fixed thresholds applied uniformly across datasets. These thresholds define a predefined operational classification and were not optimized based on the data.

Sample-level summaries were generated by aggregating interaction-level metrics within each sample, including the fraction of interactions assigned to each category and distributions of retention values.

Because classification is performed under fixed thresholds within a deterministic pipeline, regime analysis evaluates the stability of a predefined categorization rather than optimizing for separation between groups.

### Reproducibility and implementation

All analyses were implemented in Jupyter notebooks with fixed parameters and controlled stochastic components. Each dataset was processed independently under an identical pipeline, and all figures were generated from standardized outputs without manual intervention.

Because the computational framework is held constant across datasets, observed differences between samples reflect biological variation rather than analytical variability. The pipeline is modular, with separate components for preprocessing, interface scoring, null evaluation, localization, and summarization, enabling transparent inspection and reproducible execution of each stage.

### Datasets

Spatial transcriptomics datasets from breast cancer, colorectal cancer, melanoma, and pancreatic ductal adenocarcinoma were analyzed. Public datasets were obtained from the Gene Expression Omnibus (GEO) under accession numbers GSE217414, GSE243022, GSE250636, and GSE274103, along with publicly available 10x Genomics Visium datasets.

All datasets include spot-level gene expression measurements with associated spatial coordinates and were processed using the same standardized pipeline without dataset-specific modification.

## Supporting information

Supplementary Notebook 1-5

## Supplementary Information

The following Jupyter notebooks reproduce the analyses presented in this study under a locked and reproducible pipeline. Each notebook implements the same analytical framework, including interface scoring, label-permutation null evaluation, geometry-aware refinement, and sample-level summarization.

### Supplementary Notebook 1

#### Breast_cancer.ipynb

Full analysis pipeline for breast cancer spatial transcriptomics data.

### Supplementary Notebook 2

#### CRC.ipynb

Full analysis pipeline for colorectal cancer spatial transcriptomics data.

### Supplementary Notebook 3

#### Melanoma.ipynb

Full analysis pipeline for melanoma spatial transcriptomics data.

### Supplementary Notebook 4

#### PDAC.ipynb

Full analysis pipeline for pancreatic ductal adenocarcinoma spatial transcriptomics data.

### Supplementary Notebook 5

#### Figures.ipynb

Generation of figures and visualizations used in the manuscript.

