## Supplementary Notebook 1-5 for "Geometry-aware ligand-receptor analysis distinguishes interface association from spatial localization and reveals a continuum of tumor communication": Breast_cancer.ipynb

```
# %% [markdown]
# Supplementary notebook: Breast cancer locked replication pipeline

## Purpose
This notebook reproduces the locked analysis pipeline for the breast cancer dataset used in the manuscript.

## What this notebook contains
- sample setup and data loading
- preprocessing, clustering, and spatial graph construction
- interface scoring and label-permutation null evaluation
- geometry-aware localization and retention analysis
- sample-level summaries and export tables

## Notes
Code cells are preserved in execution order so the notebook continues to run as originally authored. Markdown headings were added only to improve readability for supplementary submission.


# %% [markdown]
## Initialization

Import packages, reset state, and define locked settings.

# %% [code]
for _name in [
    "adata", "adata_hvg",
    "edges_leiden_df", "boundary_leiden_df", "interface_leiden_df",
    "edges_cellchat_df", "boundary_cellchat_df", "interface_cellchat_df",
    "lr_df", "lr_obs", "lr_full", "sample_summary", "combined_summary",
    "cluster_map", "null_df", "null_scores", "null_summary_df",
    "indices", "distances", "coords", "positions",
    "interface_midpoints", "interface_scale"
]:
    if _name in globals():
        del globals()[_name]

print("Cleared prior in-memory state.")

# %% [markdown]
## Initialization

Import packages, reset state, and define locked settings.

# %% [code]
import numpy as np
import pandas as pd
import scanpy as sc
import matplotlib.pyplot as plt

from pathlib import Path
from scipy import sparse
from sklearn.neighbors import NearestNeighbors
from scipy.spatial.distance import cdist
import tarfile

pd.set_option("display.max_columns", 200)
pd.set_option("display.max_rows", 200)

# Fixed analysis parameters
N_TOP_HVG = 3000
N_NEIGHBORS_CLUSTER = 15
N_PCS = 30
LEIDEN_RESOLUTION = 0.6

N_NEIGHBORS_SPATIAL = 6
MIN_INTERFACE_EDGES = 20

TOP_N_COMPARE = 20
N_PERM = 100
RANDOM_SEED = 42

np.random.seed(RANDOM_SEED)

print("=== GENERAL SETUP LOADED ===")
print("N_TOP_HVG =", N_TOP_HVG)
print("N_NEIGHBORS_CLUSTER =", N_NEIGHBORS_CLUSTER)
print("N_PCS =", N_PCS)
print("LEIDEN_RESOLUTION =", LEIDEN_RESOLUTION)
print("N_NEIGHBORS_SPATIAL =", N_NEIGHBORS_SPATIAL)
print("MIN_INTERFACE_EDGES =", MIN_INTERFACE_EDGES)
print("TOP_N_COMPARE =", TOP_N_COMPARE)
print("N_PERM =", N_PERM)
print("RANDOM_SEED =", RANDOM_SEED)

# %% [markdown]
## Sample selection

Define the sample or sample index analyzed in this run.

# %% [code]
from pathlib import Path

TISSUE = "breast"

BASE_DIR = Path("/Users/sally/Desktop/Metrics_project/Samples/GSE243022_RAW")
LR_BASE_DIR = Path("/Users/sally/Desktop/Metrics_project/LR_tables")

BREAST_SAMPLES = [
    "GSM7777520_A1",
    "GSM7777521_B1",
    "GSM7777522_C1",
    "GSM7777523_D1",
]

SAMPLE_IDX = 2

if SAMPLE_IDX < 0 or SAMPLE_IDX >= len(BREAST_SAMPLES):
    raise IndexError(f"SAMPLE_IDX={SAMPLE_IDX} out of range for {len(BREAST_SAMPLES)} samples")

sample_folder = BREAST_SAMPLES[SAMPLE_IDX]
DATA_DIR = BASE_DIR / sample_folder
SAMPLE_NAME = sample_folder
OUT_DIR = DATA_DIR / "boundary_enriched_outputs"

LR_TABLE_PATH = LR_BASE_DIR / f"{SAMPLE_NAME}_lr.csv"
R_CLUSTER_PATH = LR_BASE_DIR / f"{SAMPLE_NAME}_clusters.csv"

h5_files = sorted(DATA_DIR.glob("*.h5"))
tar_h5_files = sorted(DATA_DIR.glob("*filtered_feature_bc_matrix*.tar.gz"))

if len(h5_files) > 0:
    COUNT_FILE = h5_files[0].name
elif len(tar_h5_files) > 0:
    COUNT_FILE = None
else:
    COUNT_FILE = None

if not DATA_DIR.exists():
    raise FileNotFoundError(f"Sample folder not found: {DATA_DIR}")

if not LR_TABLE_PATH.exists():
    raise FileNotFoundError(f"LR table not found: {LR_TABLE_PATH}")

if not R_CLUSTER_PATH.exists():
    raise FileNotFoundError(f"Cluster table not found: {R_CLUSTER_PATH}")

OUT_DIR.mkdir(parents=True, exist_ok=True)

print("=== AUTOMATIC SAMPLE SELECTION ===")
print("SAMPLE_IDX:", SAMPLE_IDX)
print("sample_folder:", sample_folder)
print("SAMPLE_NAME:", SAMPLE_NAME)
print("TISSUE:", TISSUE)
print("DATA_DIR:", DATA_DIR)
print("COUNT_FILE:", COUNT_FILE)
print("LR_TABLE_PATH:", LR_TABLE_PATH)
print("R_CLUSTER_PATH:", R_CLUSTER_PATH)
print("OUT_DIR:", OUT_DIR)

print("\nAvailable BREAST samples:")
for i, s in enumerate(BREAST_SAMPLES):
    marker = " <--- selected" if i == SAMPLE_IDX else ""
    print(f"[{i}] {s}{marker}")

# %% [markdown]
## Data loading

Load the spatial transcriptomics sample and associated metadata.

# %% [code]
import tarfile

# --------------------------------------------------
# Resolve count matrix source
# --------------------------------------------------
if COUNT_FILE is not None:
    count_path = DATA_DIR / COUNT_FILE
    if not count_path.exists():
        raise FileNotFoundError(f"Count file not found: {count_path}")
else:
    tar_candidates = sorted(DATA_DIR.glob("*filtered_feature_bc_matrix*.tar.gz"))
    if len(tar_candidates) == 0:
        raise FileNotFoundError(
            f"No .h5 file and no filtered_feature_bc_matrix tar.gz found in {DATA_DIR}"
        )

    tar_path = tar_candidates[0]
    extract_dir = DATA_DIR / "filtered_feature_bc_matrix_extracted"
    extract_dir.mkdir(parents=True, exist_ok=True)

    with tarfile.open(tar_path, "r:gz") as tar:
        tar.extractall(path=extract_dir)

    h5_candidates = sorted(extract_dir.rglob("*.h5"))
    if len(h5_candidates) == 0:
        raise FileNotFoundError(f"No .h5 file found after extracting {tar_path}")

    count_path = h5_candidates[0]

print("=== LOADING SAMPLE ===")
print("count_path:", count_path)

# --------------------------------------------------
# Read 10x Visium / feature matrix
# --------------------------------------------------
adata = sc.read_10x_h5(count_path)
adata.var_names_make_unique()

print("\nRaw AnnData shape:", adata.shape)

# --------------------------------------------------
# Attach spatial coordinates from tissue_positions
# --------------------------------------------------
positions_candidates = (
    sorted(DATA_DIR.rglob("*tissue_positions*.csv")) +
    sorted(DATA_DIR.rglob("*tissue_positions*.parquet")) +
    sorted(DATA_DIR.rglob("*tissue_positions_list*.csv"))
)

if len(positions_candidates) == 0:
    raise FileNotFoundError(f"No tissue_positions file found under {DATA_DIR}")

positions_path = positions_candidates[0]
print("positions_path:", positions_path)

if positions_path.suffix == ".parquet":
    positions = pd.read_parquet(positions_path)
else:
    try:
        positions = pd.read_csv(positions_path)
    except Exception:
        positions = pd.read_csv(positions_path, header=None)

# Handle standard 10x formats
if positions.shape[1] >= 6 and "barcode" not in positions.columns:
    positions = positions.iloc[:, :6].copy()
    positions.columns = [
        "barcode",
        "in_tissue",
        "array_row",
        "array_col",
        "pxl_row_in_fullres",
        "pxl_col_in_fullres",
    ]

required_pos_cols = {"barcode", "pxl_row_in_fullres", "pxl_col_in_fullres"}
if not required_pos_cols.issubset(set(positions.columns)):
    raise ValueError(
        f"Positions file missing required columns. Found: {positions.columns.tolist()}"
    )

positions["barcode"] = positions["barcode"].astype(str)
positions = positions.drop_duplicates(subset="barcode").set_index("barcode")

common_barcodes = adata.obs_names.intersection(positions.index)
if len(common_barcodes) == 0:
    raise RuntimeError("No overlapping barcodes between count matrix and positions file")

adata = adata[common_barcodes].copy()
positions = positions.loc[common_barcodes].copy()

adata.obsm["spatial"] = positions[["pxl_col_in_fullres", "pxl_row_in_fullres"]].to_numpy()

print("Matched spots with spatial coordinates:", adata.n_obs)
print("Genes:", adata.n_vars)
print("Spatial matrix shape:", adata.obsm["spatial"].shape)

# %% [markdown]
## Preprocessing

Perform quality control, normalization, and feature selection.

# %% [code]
adata.var_names_make_unique()

print("=== PREPROCESSING ===")
print("Input shape:", adata.shape)

# ---------- QC metrics ----------
if sparse.issparse(adata.X):
    n_counts = np.asarray(adata.X.sum(axis=1)).ravel()
    n_genes = np.asarray((adata.X > 0).sum(axis=1)).ravel()
else:
    n_counts = adata.X.sum(axis=1).ravel()
    n_genes = (adata.X > 0).sum(axis=1).ravel()

adata.obs["n_counts"] = n_counts
adata.obs["n_genes"] = n_genes

adata.var["mt"] = adata.var_names.str.upper().str.startswith("MT-")
mt_mask = adata.var["mt"].to_numpy()

if mt_mask.sum() > 0:
    if sparse.issparse(adata.X):
        mt_counts = np.asarray(adata[:, mt_mask].X.sum(axis=1)).ravel()
    else:
        mt_counts = adata[:, mt_mask].X.sum(axis=1).ravel()
    adata.obs["pct_counts_mt"] = 100 * mt_counts / np.maximum(adata.obs["n_counts"].to_numpy(), 1)
else:
    adata.obs["pct_counts_mt"] = 0.0

print("\nQC summary before filtering:")
display(adata.obs[["n_counts", "n_genes", "pct_counts_mt"]].describe().T)

# ---------- Locked minimal filtering ----------
sc.pp.filter_cells(adata, min_genes=200)
sc.pp.filter_genes(adata, min_cells=3)

print("\nShape after minimal filtering:", adata.shape)

# ---------- Store raw counts before normalization ----------
adata.layers["counts"] = adata.X.copy()

# ---------- Normalize and log-transform ----------
sc.pp.normalize_total(adata, target_sum=1e4)
sc.pp.log1p(adata)

# ---------- HVGs ----------
sc.pp.highly_variable_genes(
    adata,
    flavor="seurat",
    n_top_genes=N_TOP_HVG
)

n_hvg = int(adata.var["highly_variable"].sum())
print("Number of HVGs selected:", n_hvg)

if n_hvg == 0:
    raise RuntimeError("No highly variable genes were selected.")

print("\nExample HVGs:")
display(adata.var.loc[adata.var["highly_variable"]].head(10))

print("\nFinal shape after preprocessing:", adata.shape)

# %% [markdown]
## Dimensionality reduction and clustering

Compute PCA, neighborhood graph, embedding, and Leiden clusters under the locked pipeline.

# %% [code]
print("=== PCA + CLUSTERING ===")
print("Input shape:", adata.shape)

# Use HVGs for dimensionality reduction and clustering
adata_hvg = adata[:, adata.var["highly_variable"]].copy()

# Scaling for PCA/clustering
sc.pp.scale(adata_hvg, max_value=10)

# PCA
sc.tl.pca(adata_hvg, n_comps=N_PCS, svd_solver="arpack", random_state=RANDOM_SEED)

# Neighbor graph in expression space
sc.pp.neighbors(
    adata_hvg,
    n_neighbors=N_NEIGHBORS_CLUSTER,
    n_pcs=N_PCS,
    random_state=RANDOM_SEED
)

# UMAP
sc.tl.umap(adata_hvg, random_state=RANDOM_SEED)

# Leiden clustering
sc.tl.leiden(
    adata_hvg,
    resolution=LEIDEN_RESOLUTION,
    key_added="leiden",
    random_state=RANDOM_SEED
)

# Transfer embeddings and clusters back to full adata
adata.obsm["X_pca"] = adata_hvg.obsm["X_pca"].copy()
adata.obsm["X_umap"] = adata_hvg.obsm["X_umap"].copy()
adata.obs["leiden"] = adata_hvg.obs["leiden"].astype(str).copy()

print("PCA shape:", adata.obsm["X_pca"].shape)
print("UMAP shape:", adata.obsm["X_umap"].shape)
print("Number of Leiden clusters:", adata.obs["leiden"].nunique())

print("\nCluster sizes:")
display(
    adata.obs["leiden"]
    .value_counts()
    .sort_index()
    .rename_axis("cluster")
    .reset_index(name="n_spots")
)

print("\nExplained variance ratio (first 10 PCs):")
print(np.round(adata_hvg.uns["pca"]["variance_ratio"][:10], 4))

# %% [markdown]
## Cluster inspection

Visualize clusters and verify locked spatial metadata.

# %% [code]
print("=== CLUSTER VISUALIZATION ===")

if "leiden" not in adata.obs.columns:
    raise RuntimeError("adata.obs['leiden'] is missing.")
if "X_umap" not in adata.obsm:
    raise RuntimeError("adata.obsm['X_umap'] is missing.")
if "spatial" not in adata.obsm:
    raise RuntimeError("adata.obsm['spatial'] is missing.")

print("Leiden clusters:", sorted(adata.obs["leiden"].astype(str).unique().tolist()))
print("UMAP shape:", adata.obsm["X_umap"].shape)
print("Spatial shape:", adata.obsm["spatial"].shape)

fig, axes = plt.subplots(1, 2, figsize=(12, 5))

# UMAP
sc.pl.umap(
    adata,
    color="leiden",
    ax=axes[0],
    show=False,
    title="UMAP — Leiden clusters",
    legend_loc="on data"
)

# Spatial
spatial_df = pd.DataFrame(
    adata.obsm["spatial"],
    index=adata.obs_names,
    columns=["x", "y"]
).join(adata.obs["leiden"])

for cl in sorted(spatial_df["leiden"].astype(str).unique()):
    sub = spatial_df[spatial_df["leiden"].astype(str) == cl]
    axes[1].scatter(
        sub["x"],
        sub["y"],
        s=8,
        label=str(cl)
    )

axes[1].invert_yaxis()
axes[1].set_title("Spatial coordinates — Leiden clusters")
axes[1].set_xlabel("x")
axes[1].set_ylabel("y")
axes[1].legend(
    title="leiden",
    bbox_to_anchor=(1.05, 1),
    loc="upper left",
    markerscale=2
)

plt.tight_layout()
plt.show()

# %% [markdown]
## Spatial graph construction

Build the spatial nearest-neighbor graph used for interface analysis.

# %% [code]
print("=== SPATIAL kNN GRAPH ===")

if "spatial" not in adata.obsm:
    raise RuntimeError("adata.obsm['spatial'] is missing.")

coords = np.asarray(adata.obsm["spatial"], dtype=float)

if coords.ndim != 2 or coords.shape[1] != 2:
    raise RuntimeError(f"Expected spatial coordinates with shape (n_spots, 2), got {coords.shape}")

n_spots = coords.shape[0]
if N_NEIGHBORS_SPATIAL >= n_spots:
    raise RuntimeError(
        f"N_NEIGHBORS_SPATIAL={N_NEIGHBORS_SPATIAL} must be smaller than number of spots={n_spots}"
    )

# +1 because the nearest neighbor is the point itself
knn = NearestNeighbors(n_neighbors=N_NEIGHBORS_SPATIAL + 1, metric="euclidean")
knn.fit(coords)

distances_full, indices_full = knn.kneighbors(coords)

# remove self-neighbor in column 0
distances = distances_full[:, 1:].copy()
indices = indices_full[:, 1:].copy()

print("Number of spots:", n_spots)
print("Spatial neighbors per spot:", N_NEIGHBORS_SPATIAL)
print("indices shape:", indices.shape)
print("distances shape:", distances.shape)

flat_d = distances.ravel()
print("\nDistance summary:")
print(pd.Series(flat_d).describe())

# store for downstream cells
adata.uns["spatial_knn"] = {
    "n_neighbors": N_NEIGHBORS_SPATIAL,
    "indices_shape": indices.shape,
    "distances_shape": distances.shape,
}

# %% [markdown]
## Leiden-based interfaces

Construct interface tables using Leiden cluster labels.

# %% [code]
print("=== LEIDEN-BASED INTERFACE TABLE ===")

if "leiden" not in adata.obs.columns:
    raise RuntimeError("adata.obs['leiden'] is missing.")
if "spatial" not in adata.obsm:
    raise RuntimeError("adata.obsm['spatial'] is missing.")
if "indices" not in globals() or "distances" not in globals():
    raise RuntimeError("indices/distances missing. Run Cell 7 first.")

labels = adata.obs["leiden"].astype(str).to_numpy()
spot_ids = adata.obs_names.to_numpy()

edge_rows = []
for i in range(indices.shape[0]):
    for j_pos, j in enumerate(indices[i]):
        j = int(j)

        # keep each undirected edge once
        if i >= j:
            continue

        li = labels[i]
        lj = labels[j]
        d = float(distances[i, j_pos])

        if d <= 0:
            continue

        edge_rows.append(
            {
                "spot_i": spot_ids[i],
                "spot_j": spot_ids[j],
                "idx_i": i,
                "idx_j": j,
                "source": li,
                "target": lj,
                "cluster_pair": "|".join(sorted([li, lj])),
                "distance": d,
                "weight": 1.0 / (d + 1e-8),
            }
        )

edges_leiden_df = pd.DataFrame(edge_rows)

if edges_leiden_df.empty:
    raise RuntimeError("No spatial edges were created.")

edges_leiden_df["is_boundary"] = edges_leiden_df["source"] != edges_leiden_df["target"]
boundary_leiden_df = edges_leiden_df.loc[edges_leiden_df["is_boundary"]].copy()

if boundary_leiden_df.empty:
    raise RuntimeError("No Leiden boundary edges were found.")

interface_leiden_df = (
    boundary_leiden_df.groupby("cluster_pair", as_index=False)
    .agg(
        n_boundary_edges=("cluster_pair", "size"),
        boundary_weight_sum=("weight", "sum"),
        mean_edge_distance=("distance", "mean"),
    )
)

# Locked boundary definition
interface_leiden_df["boundary_score"] = interface_leiden_df["boundary_weight_sum"]

# Locked support flag
interface_leiden_df["keep_interface"] = (
    interface_leiden_df["n_boundary_edges"] >= MIN_INTERFACE_EDGES
)

interface_leiden_df = interface_leiden_df.sort_values(
    ["keep_interface", "boundary_score"],
    ascending=[False, False]
).reset_index(drop=True)

print("Total undirected edges:", len(edges_leiden_df))
print("Boundary edges:", len(boundary_leiden_df))
print("Unique Leiden interfaces:", interface_leiden_df.shape[0])
print("Interfaces kept:", int(interface_leiden_df["keep_interface"].sum()))
print("Interfaces removed:", int((~interface_leiden_df["keep_interface"]).sum()))

display(interface_leiden_df.head(15))

# %% [markdown]
## Ligand–receptor table integration

Load ligand–receptor results and align CellChat sender/receiver labels to the spatial object.

# %% [code]
print("=== LR TABLE + CELLCHAT LABELS ===")

# -------- load inputs --------
lr_df = pd.read_csv(LR_TABLE_PATH)
r_clusters = pd.read_csv(R_CLUSTER_PATH)

print("SAMPLE_NAME:", SAMPLE_NAME)
print("LR table shape:", lr_df.shape)
print("R cluster map shape:", r_clusters.shape)

# -------- validate cluster map --------
required_cluster_cols = ["barcode", "cellchat_label"]
missing_cluster_cols = [c for c in required_cluster_cols if c not in r_clusters.columns]
if len(missing_cluster_cols) > 0:
    raise ValueError(f"Missing required columns in cluster file: {missing_cluster_cols}")

cluster_map = r_clusters[["barcode", "cellchat_label"]].copy()
cluster_map["barcode"] = cluster_map["barcode"].astype(str)
cluster_map["cellchat_label"] = cluster_map["cellchat_label"].astype(str)

# remove exact duplicate barcode rows if present
cluster_map = cluster_map.drop_duplicates(subset="barcode", keep="first").copy()

# -------- attach labels to AnnData WITHOUT row expansion --------
adata.obs = adata.obs.copy()
adata.obs["barcode"] = adata.obs_names.astype(str)

barcode_to_label = dict(zip(cluster_map["barcode"], cluster_map["cellchat_label"]))
adata.obs["cellchat_label"] = adata.obs["barcode"].map(barcode_to_label)

missing_labels = int(adata.obs["cellchat_label"].isna().sum())
matched_labels = int(adata.obs["cellchat_label"].notna().sum())

print("\nMatched CellChat labels:", matched_labels)
print("Missing CellChat labels:", missing_labels)

if matched_labels == 0:
    raise RuntimeError("No CellChat labels matched adata.obs_names. Barcode mapping failed.")

print("\nCellChat label counts:")
display(
    adata.obs["cellchat_label"]
    .value_counts(dropna=False)
    .sort_index()
    .rename_axis("cellchat_label")
    .reset_index(name="n_spots")
)

# -------- validate LR table --------
required_lr_cols = [
    "source", "target", "ligand", "receptor",
    "prob", "pval", "interaction_name", "pathway_name"
]
missing_lr_cols = [c for c in required_lr_cols if c not in lr_df.columns]
if len(missing_lr_cols) > 0:
    raise ValueError(f"Missing required columns in LR table: {missing_lr_cols}")

lr_df = lr_df[required_lr_cols].copy()

for col in ["source", "target", "ligand", "receptor", "interaction_name", "pathway_name"]:
    lr_df[col] = lr_df[col].astype(str)

lr_df["prob"] = pd.to_numeric(lr_df["prob"], errors="coerce")
lr_df["pval"] = pd.to_numeric(lr_df["pval"], errors="coerce")

lr_df = lr_df.dropna(subset=["source", "target", "ligand", "receptor", "prob", "pval"]).copy()
lr_df = lr_df[lr_df["pval"] <= 0.05].copy()

valid_labels = set(adata.obs["cellchat_label"].dropna().astype(str).unique())
lr_df = lr_df[
    lr_df["source"].isin(valid_labels) &
    lr_df["target"].isin(valid_labels)
].copy()

if lr_df.empty:
    raise RuntimeError("No significant LR interactions remain after restricting to observed CellChat labels.")

lr_df = lr_df.sort_values(["prob", "interaction_name"], ascending=[False, True]).reset_index(drop=True)

print("\nSignificant LR interactions after label restriction:", lr_df.shape[0])
print("Unique LR source labels:", lr_df["source"].nunique())
print("Unique LR target labels:", lr_df["target"].nunique())

print("\nTop LR rows:")
display(lr_df.head(15))

# %% [markdown]
## CellChat-based interfaces

Recompute supported interfaces using CellChat-derived labels.

# %% [code]
print("=== CELLCHAT-BASED INTERFACE TABLE ===")

# Required objects from previous cells
if "adata" not in globals():
    raise RuntimeError("adata is missing. Run previous cells first.")
if "indices" not in globals() or "distances" not in globals():
    raise RuntimeError("indices/distances are missing. Run Cell 7 first.")
if "cellchat_label" not in adata.obs.columns:
    raise RuntimeError("adata.obs['cellchat_label'] is missing. Run Cell 9 first.")

cellchat_labels = adata.obs["cellchat_label"].astype(str).to_numpy()
spot_ids = adata.obs_names.to_numpy()

edge_rows = []
for i in range(indices.shape[0]):
    for j_pos, j in enumerate(indices[i]):
        j = int(j)

        # keep each undirected edge only once
        if i >= j:
            continue

        li = cellchat_labels[i]
        lj = cellchat_labels[j]
        d = float(distances[i, j_pos])

        if pd.isna(li) or pd.isna(lj):
            continue
        if d <= 0:
            continue

        edge_rows.append(
            {
                "spot_i": spot_ids[i],
                "spot_j": spot_ids[j],
                "idx_i": i,
                "idx_j": j,
                "source": li,
                "target": lj,
                "cluster_pair": "|".join(sorted([li, lj])),
                "distance": d,
                "weight": 1.0 / (d + 1e-8),
            }
        )

edges_cellchat_df = pd.DataFrame(edge_rows)

if edges_cellchat_df.empty:
    raise RuntimeError("No spatial edges were created with CellChat labels.")

edges_cellchat_df["is_boundary"] = edges_cellchat_df["source"] != edges_cellchat_df["target"]
boundary_cellchat_df = edges_cellchat_df.loc[edges_cellchat_df["is_boundary"]].copy()

if boundary_cellchat_df.empty:
    raise RuntimeError("No CellChat boundary edges were found.")

interface_cellchat_df = (
    boundary_cellchat_df.groupby("cluster_pair", as_index=False)
    .agg(
        n_boundary_edges=("cluster_pair", "size"),
        boundary_weight_sum=("weight", "sum"),
        mean_edge_distance=("distance", "mean"),
    )
)

# Locked definition: boundary score = sum of inverse-distance boundary weights
interface_cellchat_df["boundary_score"] = interface_cellchat_df["boundary_weight_sum"]

# Locked interface support filter
interface_cellchat_df["keep_interface"] = (
    interface_cellchat_df["n_boundary_edges"] >= MIN_INTERFACE_EDGES
)

interface_cellchat_df = interface_cellchat_df.sort_values(
    ["keep_interface", "boundary_score"],
    ascending=[False, False]
).reset_index(drop=True)

print("Total undirected edges:", len(edges_cellchat_df))
print("Boundary edges:", len(boundary_cellchat_df))
print("Unique CellChat interfaces:", interface_cellchat_df.shape[0])
print("Interfaces kept:", int(interface_cellchat_df["keep_interface"].sum()))
print("Interfaces removed:", int((~interface_cellchat_df["keep_interface"]).sum()))

print("\nTop CellChat interfaces:")
display(interface_cellchat_df.head(15))

# %% [markdown]
## Observed ligand–receptor mapping

Map observed ligand–receptor interactions to supported interfaces.

# %% [code]
print("=== OBSERVED LR-INTERFACE MAPPING ===")

if "lr_df" not in globals():
    raise RuntimeError("lr_df is missing. Run Cell 9 first.")
if "interface_cellchat_df" not in globals():
    raise RuntimeError("interface_cellchat_df is missing. Run Cell 10 first.")

# Keep only supported interfaces
supported_interfaces = interface_cellchat_df.loc[
    interface_cellchat_df["keep_interface"]
].copy()

if supported_interfaces.empty:
    raise RuntimeError("No supported CellChat interfaces remain after MIN_INTERFACE_EDGES filtering.")

interface_lookup = supported_interfaces[
    ["cluster_pair", "n_boundary_edges", "boundary_score", "mean_edge_distance"]
].copy()

# Direction-aware sender/receiver retained; spatial interface is undirected
lr_obs = lr_df.copy()
lr_obs["sender_cluster"] = lr_obs["source"].astype(str)
lr_obs["receiver_cluster"] = lr_obs["target"].astype(str)
lr_obs["cluster_pair"] = lr_obs.apply(
    lambda r: "|".join(sorted([r["sender_cluster"], r["receiver_cluster"]])),
    axis=1
)

lr_obs = lr_obs.merge(
    interface_lookup,
    how="left",
    on="cluster_pair",
    validate="many_to_one"
)

n_total = lr_obs.shape[0]
n_matched = int(lr_obs["boundary_score"].notna().sum())
n_unmatched = int(lr_obs["boundary_score"].isna().sum())

print("Total significant LR rows:", n_total)
print("Matched to supported interfaces:", n_matched)
print("Unmatched to supported interfaces:", n_unmatched)

# Keep only LR interactions assigned to supported interfaces
lr_obs = lr_obs.dropna(subset=["boundary_score"]).copy()

if lr_obs.empty:
    raise RuntimeError("No LR interactions mapped to supported CellChat interfaces.")

# Standard locked column names
lr_obs = lr_obs.rename(
    columns={
        "prob": "cellchat_prob",
        "pval": "cellchat_pval",
    }
)

lr_obs = lr_obs[
    [
        "source",
        "target",
        "sender_cluster",
        "receiver_cluster",
        "cluster_pair",
        "ligand",
        "receptor",
        "interaction_name",
        "pathway_name",
        "cellchat_prob",
        "cellchat_pval",
        "n_boundary_edges",
        "mean_edge_distance",
        "boundary_score",
    ]
].copy()

lr_obs = lr_obs.sort_values(
    ["boundary_score", "cellchat_prob", "interaction_name"],
    ascending=[False, False, True]
).reset_index(drop=True)

print("\nObserved LR rows after interface mapping:", lr_obs.shape[0])
print("Unique mapped interfaces:", lr_obs["cluster_pair"].nunique())
print("Unique pathways:", lr_obs["pathway_name"].nunique())

print("\nTop mapped LR rows:")
display(lr_obs.head(20))

# %% [markdown]
## Label-permutation null model

Compute interface null distributions by permuting labels on fixed geometry.

# %% [code]
print("=== NULL MODEL (LABEL PERMUTATION) ===")

if "interface_cellchat_df" not in globals():
    raise RuntimeError("interface_cellchat_df missing.")
if "boundary_cellchat_df" not in globals():
    raise RuntimeError("boundary_cellchat_df missing.")
if "lr_obs" not in globals():
    raise RuntimeError("lr_obs missing.")

# Precompute static edge structure (indices + distances already fixed)
edge_index_i = boundary_cellchat_df["idx_i"].to_numpy()
edge_index_j = boundary_cellchat_df["idx_j"].to_numpy()
edge_distances = boundary_cellchat_df["distance"].to_numpy()

n_edges = len(edge_index_i)
print("Boundary edges used for null:", n_edges)

# Observed labels
labels_obs = adata.obs["cellchat_label"].astype(str).to_numpy()
unique_labels = np.unique(labels_obs)

# Store null scores per cluster_pair
cluster_pairs = interface_cellchat_df["cluster_pair"].tolist()
null_scores = {cp: [] for cp in cluster_pairs}

rng = np.random.default_rng(RANDOM_SEED)

for perm in range(N_PERM):

    permuted_labels = rng.permutation(labels_obs)

    pair_weights = {}

    for k in range(n_edges):
        i = edge_index_i[k]
        j = edge_index_j[k]

        li = permuted_labels[i]
        lj = permuted_labels[j]

        if li == lj:
            continue

        cp = "|".join(sorted([li, lj]))

        w = 1.0 / (edge_distances[k] + 1e-8)

        if cp not in pair_weights:
            pair_weights[cp] = 0.0

        pair_weights[cp] += w

    # record results for all cluster pairs (including missing → 0)
    for cp in cluster_pairs:
        null_scores[cp].append(pair_weights.get(cp, 0.0))

    if (perm + 1) % 10 == 0:
        print(f"Permutation {perm+1}/{N_PERM}")

# convert to DataFrame
null_df = pd.DataFrame({
    "cluster_pair": cluster_pairs,
    "null_mean": [np.mean(null_scores[cp]) for cp in cluster_pairs]
})

print("\nNull model completed")
display(null_df.head())

# %% [markdown]
## Null merge and empirical p-values

Merge null results back onto observed interfaces and compute empirical support.

# %% [code]
print("=== LR + NULL MERGE ===")

if "lr_obs" not in globals():
    raise RuntimeError("lr_obs missing.")
if "null_scores" not in globals():
    raise RuntimeError("null_scores missing.")

# Convert null distributions to array form
cluster_pairs = list(null_scores.keys())

null_summary_rows = []

for cp in cluster_pairs:
    values = np.array(null_scores[cp])
    null_summary_rows.append({
        "cluster_pair": cp,
        "null_mean": values.mean(),
        "null_values": values  # keep full distribution for p-values
    })

null_summary_df = pd.DataFrame(null_summary_rows)

# Merge into LR table
lr_full = lr_obs.merge(
    null_summary_df,
    on="cluster_pair",
    how="left",
    validate="many_to_one"
)

if lr_full["null_mean"].isna().any():
    raise RuntimeError("Null values missing after merge.")

# -------- empirical p-values --------
p_values = []

for _, row in lr_full.iterrows():
    B = row["boundary_score"]
    null_dist = row["null_values"]

    count = np.sum(null_dist >= B)
    p = (1 + count) / (len(null_dist) + 1)

    p_values.append(p)

lr_full["empirical_p"] = p_values

# -------- derived quantities --------
lr_full["null_gap"] = lr_full["boundary_score"] - lr_full["null_mean"]

lr_full["null_gap_fraction"] = (
    lr_full["null_gap"] / lr_full["null_mean"].replace(0, np.nan)
)

# clean structure
lr_full = lr_full.drop(columns=["null_values"])

print("LR rows:", lr_full.shape[0])
print("Mean empirical p:", lr_full["empirical_p"].mean())

print("\nPreview:")
display(lr_full.head(15))

# %% [markdown]
## Geometry-aware refinement

Compute ligand–receptor localization relative to interfaces and derive geometry-aware scores.

# %% [code]
print("=== LR-SPECIFIC LOCALIZATION + GEOMETRY SCORE ===")

if "boundary_cellchat_df" not in globals():
    raise RuntimeError("boundary_cellchat_df missing.")
if "lr_full" not in globals():
    raise RuntimeError("lr_full missing.")
if "adata" not in globals():
    raise RuntimeError("adata missing.")

coords = np.asarray(adata.obsm["spatial"], dtype=float)
spot_labels = adata.obs["cellchat_label"].astype(str).to_numpy()
gene_index = pd.Index(adata.var_names.astype(str))

gene_cache = {}
complex_cache = {}
cluster_index_cache = {}

def get_gene_vector(gene_name):
    gene_name = str(gene_name)
    if gene_name in gene_cache:
        return gene_cache[gene_name]
    if gene_name in gene_index:
        x = adata[:, [gene_name]].X
        if sparse.issparse(x):
            v = np.asarray(x.toarray()).ravel()
        else:
            v = np.asarray(x).ravel()
        gene_cache[gene_name] = v
        return v
    gene_cache[gene_name] = None
    return None

def get_complex_vector(feature_name):
    feature_name = str(feature_name)
    if feature_name in complex_cache:
        return complex_cache[feature_name]
    parts = [p for p in feature_name.split("_") if p != ""]
    vecs = []
    for p in parts:
        v = get_gene_vector(p)
        if v is not None:
            vecs.append(v)
    if len(vecs) == 0:
        complex_cache[feature_name] = None
        return None
    if len(vecs) == 1:
        complex_cache[feature_name] = vecs[0]
        return vecs[0]
    out = np.vstack(vecs).min(axis=0)
    complex_cache[feature_name] = out
    return out

# interface geometry
interface_midpoints = {}
interface_scale = {}

for cp, sub in boundary_cellchat_df.groupby("cluster_pair"):
    i_idx = sub["idx_i"].to_numpy()
    j_idx = sub["idx_j"].to_numpy()

    mids = (coords[i_idx] + coords[j_idx]) / 2.0
    interface_midpoints[cp] = mids

    scale = float(np.median(sub["distance"].to_numpy()))
    if not np.isfinite(scale) or scale <= 0:
        scale = 1.0
    interface_scale[cp] = scale

print("Computed interface geometry for:", len(interface_midpoints), "interfaces")

for label in np.unique(spot_labels):
    cluster_index_cache[str(label)] = np.where(spot_labels == str(label))[0]

localization_scores = []
missing_feature_rows = 0

for row in lr_full.itertuples(index=False):
    sender = str(row.sender_cluster)
    receiver = str(row.receiver_cluster)
    cp = str(row.cluster_pair)
    ligand = str(row.ligand)
    receptor = str(row.receptor)

    if cp not in interface_midpoints:
        localization_scores.append(np.nan)
        continue

    mids = interface_midpoints[cp]
    scale = interface_scale[cp]

    sender_idx = cluster_index_cache.get(sender, np.array([], dtype=int))
    receiver_idx = cluster_index_cache.get(receiver, np.array([], dtype=int))

    if len(sender_idx) == 0 or len(receiver_idx) == 0:
        localization_scores.append(np.nan)
        continue

    ligand_vec = get_complex_vector(ligand)
    receptor_vec = get_complex_vector(receptor)

    if ligand_vec is None or receptor_vec is None:
        localization_scores.append(np.nan)
        missing_feature_rows += 1
        continue

    ligand_sender = ligand_vec[sender_idx]
    receptor_receiver = receptor_vec[receiver_idx]

    ligand_mask = ligand_sender > 0
    receptor_mask = receptor_receiver > 0

    if ligand_mask.sum() == 0 or receptor_mask.sum() == 0:
        localization_scores.append(0.0)
        continue

    sender_coords = coords[sender_idx][ligand_mask]
    receiver_coords = coords[receiver_idx][receptor_mask]

    ligand_w = ligand_sender[ligand_mask].astype(float)
    receptor_w = receptor_receiver[receptor_mask].astype(float)

    d_sender = cdist(sender_coords, mids).min(axis=1)
    d_receiver = cdist(receiver_coords, mids).min(axis=1)

    s_sender = np.exp(-d_sender / scale)
    s_receiver = np.exp(-d_receiver / scale)

    sender_score = float(np.average(s_sender, weights=ligand_w))
    receiver_score = float(np.average(s_receiver, weights=receptor_w))

    localization = 0.5 * (sender_score + receiver_score)
    localization_scores.append(localization)

lr_full["localization_score"] = localization_scores

n_nan = int(lr_full["localization_score"].isna().sum())
print("Rows with missing localization:", n_nan)
print("Rows with missing ligand/receptor features:", missing_feature_rows)

lr_full = lr_full.dropna(subset=["localization_score"]).copy()

if lr_full.empty:
    raise RuntimeError("All LR rows were removed after localization calculation.")

lr_full["geometry_score"] = lr_full["boundary_score"] * lr_full["localization_score"]

print("LR rows retained:", lr_full.shape[0])
print("Mean localization score:", float(lr_full["localization_score"].mean()))
print("Mean geometry-aware score:", float(lr_full["geometry_score"].mean()))

print("\nPreview:")
display(
    lr_full[
        [
            "sender_cluster", "receiver_cluster", "ligand", "receptor",
            "cluster_pair", "boundary_score", "null_mean", "empirical_p",
            "localization_score", "geometry_score"
        ]
    ].head(20)
)

# %% [markdown]
## Sample-level summary

Summarize retention, null-related metrics, and regime assignments for the analyzed sample.

# %% [code]
print("=== SAMPLE SUMMARY + REGIME ===")

if "lr_full" not in globals():
    raise RuntimeError("lr_full missing.")

lr_full = lr_full.copy()

lr_full["retention"] = (
    lr_full["geometry_score"] / lr_full["boundary_score"].replace(0, np.nan)
)

lr_full["combined_score"] = lr_full["geometry_score"]

lr_full["interaction_regime"] = "mixed"
lr_full.loc[
    (lr_full["retention"] >= 0.75) &
    (lr_full["null_gap_fraction"] > -0.10),
    "interaction_regime"
] = "localized"
lr_full.loc[
    (lr_full["retention"] <= 0.10) &
    (lr_full["empirical_p"] >= 0.05),
    "interaction_regime"
] = "diffuse"

lr_collapsed = (
    lr_full
    .groupby(["cluster_pair", "interaction_name"], as_index=False)
    .agg(
        boundary_score=("boundary_score", "mean"),
        null_mean=("null_mean", "mean"),
        empirical_p=("empirical_p", "mean"),
        geometry_score=("geometry_score", "mean"),
        combined_score=("combined_score", "mean"),
        retention=("retention", "mean"),
        null_gap=("null_gap", "mean"),
        null_gap_fraction=("null_gap_fraction", "mean"),
        interaction_regime=("interaction_regime", lambda x: x.mode().iloc[0]),
    )
)

print("LR rows before collapse:", lr_full.shape[0])
print("Collapsed LR rows:", lr_collapsed.shape[0])

sample_summary = pd.DataFrame([{
    "sample_id": SAMPLE_NAME,
    "tissue": TISSUE,
    "n_lr": int(lr_collapsed.shape[0]),
    "mean_boundary": float(lr_collapsed["boundary_score"].mean()),
    "mean_null": float(lr_collapsed["null_mean"].mean()),
    "mean_p": float(lr_collapsed["empirical_p"].mean()),
    "mean_geometry": float(lr_collapsed["geometry_score"].mean()),
    "mean_combined": float(lr_collapsed["combined_score"].mean()),
    "retention": float(lr_collapsed["retention"].mean()),
    "null_gap": float(lr_collapsed["null_gap"].mean()),
    "null_gap_fraction": float(lr_collapsed["null_gap_fraction"].mean()),
    "frac_localized": float((lr_collapsed["interaction_regime"] == "localized").mean()),
    "frac_mixed": float((lr_collapsed["interaction_regime"] == "mixed").mean()),
    "frac_diffuse": float((lr_collapsed["interaction_regime"] == "diffuse").mean()),
}])

sample_summary["regime"] = "mixed"
sample_summary.loc[
    (sample_summary["retention"] >= 0.75) &
    (sample_summary["null_gap_fraction"] > -0.10),
    "regime"
] = "localized"
sample_summary.loc[
    (sample_summary["retention"] <= 0.10) &
    (sample_summary["mean_p"] >= 0.05),
    "regime"
] = "diffuse"

print("\nInteraction regime counts (row-level):")
display(
    lr_full["interaction_regime"]
    .value_counts()
    .rename_axis("interaction_regime")
    .reset_index(name="n_lr_rows")
)

print("\nInteraction regime counts (collapsed):")
display(
    lr_collapsed["interaction_regime"]
    .value_counts()
    .rename_axis("interaction_regime")
    .reset_index(name="n_lr_collapsed")
)

print("\nSample summary:")
display(sample_summary)

print("\nPreview LR row table:")
display(
    lr_full[
        [
            "sender_cluster", "receiver_cluster", "cluster_pair",
            "ligand", "receptor", "interaction_name", "pathway_name",
            "boundary_score", "null_mean", "empirical_p",
            "localization_score", "geometry_score", "combined_score",
            "retention", "null_gap", "null_gap_fraction",
            "interaction_regime"
        ]
    ].head(20)
)

print("\nPreview collapsed LR table:")
display(lr_collapsed.head(20))

# %% [markdown]
## Export of sample results

Write per-sample outputs and locked summary tables used downstream.

# %% [code]
print("=== SAVE CURRENT SAMPLE RESULTS ===")

if "lr_full" not in globals():
    raise RuntimeError("lr_full missing.")
if "lr_collapsed" not in globals():
    raise RuntimeError("lr_collapsed missing.")
if "sample_summary" not in globals():
    raise RuntimeError("sample_summary missing.")
if "OUT_DIR" not in globals():
    raise RuntimeError("OUT_DIR missing.")
if "BASE_DIR" not in globals():
    raise RuntimeError("BASE_DIR missing.")

lr_row_path = OUT_DIR / f"{SAMPLE_NAME}_lr_row_table.csv"
lr_collapsed_path = OUT_DIR / f"{SAMPLE_NAME}_lr_collapsed_table.csv"
sample_summary_path = OUT_DIR / f"{SAMPLE_NAME}_sample_summary.csv"

lr_full.to_csv(lr_row_path, index=False)
lr_collapsed.to_csv(lr_collapsed_path, index=False)
sample_summary.to_csv(sample_summary_path, index=False)

print("Saved LR row table:", lr_row_path)
print("Saved LR collapsed table:", lr_collapsed_path)
print("Saved sample summary:", sample_summary_path)

print("\n=== FOUND SAMPLE SUMMARY FILES ===")

summary_files = sorted(
    BASE_DIR.glob("**/boundary_enriched_outputs/*_sample_summary.csv")
)

for p in summary_files:
    print(p)

print("\nTotal sample summaries found:", len(summary_files))

if len(summary_files) == 0:
    raise RuntimeError("No sample summary files found.")

summary_list = []
for p in summary_files:
    df = pd.read_csv(p)
    df["summary_file"] = str(p)
    summary_list.append(df)

combined_summary = pd.concat(summary_list, ignore_index=True)

preferred_cols = [
    "sample_id",
    "tissue",
    "n_lr",
    "mean_boundary",
    "mean_null",
    "mean_p",
    "mean_geometry",
    "mean_combined",
    "retention",
    "null_gap",
    "null_gap_fraction",
    "frac_localized",
    "frac_mixed",
    "frac_diffuse",
    "regime",
    "summary_file",
]
existing_cols = [c for c in preferred_cols if c in combined_summary.columns]
combined_summary = combined_summary[existing_cols].copy()

combined_summary = combined_summary.sort_values(
    ["tissue", "sample_id"]
).reset_index(drop=True)

combined_out_dir = BASE_DIR / "combined_regime_summary"
combined_out_dir.mkdir(parents=True, exist_ok=True)

combined_summary_path = combined_out_dir / "breast_combined_sample_summary.csv"
combined_summary.to_csv(combined_summary_path, index=False)

print("\n=== COMBINED SUMMARY ===")
display(combined_summary)

print("\nSaved combined summary:", combined_summary_path)
print("\n=== DONE ===")

# %% [markdown]
## Quality control checks

Run consistency checks on filtered ligand–receptor outputs.

# %% [code]
print("=== LR FILTERING QC ===")

if "lr_df" not in globals():
    raise RuntimeError("lr_df missing (Cell 9 not run).")
if "adata" not in globals():
    raise RuntimeError("adata missing.")

# -----------------------------------
# Basic counts
# -----------------------------------
print("Total LR rows after filtering:", lr_df.shape[0])
print("Unique interactions:", lr_df["interaction_name"].nunique())
print("Unique pathways:", lr_df["pathway_name"].nunique())
print("Unique sources:", lr_df["source"].nunique())
print("Unique targets:", lr_df["target"].nunique())

# -----------------------------------
# P-value distribution
# -----------------------------------
print("\nP-value summary:")
display(lr_df["pval"].describe())

print("Min pval:", lr_df["pval"].min())
print("Max pval:", lr_df["pval"].max())

if not (lr_df["pval"] <= 0.05).all():
    raise RuntimeError("LR table contains p-values > 0.05 (filter inconsistency).")

# -----------------------------------
# Probability distribution
# -----------------------------------
print("\nProbability summary:")
display(lr_df["prob"].describe())

# -----------------------------------
# CellChat label consistency
# -----------------------------------
valid_labels = set(adata.obs["cellchat_label"].astype(str).unique())

invalid_sources = set(lr_df["source"]) - valid_labels
invalid_targets = set(lr_df["target"]) - valid_labels

print("\nInvalid sources:", invalid_sources)
print("Invalid targets:", invalid_targets)

if len(invalid_sources) > 0 or len(invalid_targets) > 0:
    raise RuntimeError("LR table contains labels not present in adata.")

# -----------------------------------
# Cluster pair distribution
# -----------------------------------
lr_df["cluster_pair"] = lr_df.apply(
    lambda r: "|".join(sorted([str(r["source"]), str(r["target"])])),
    axis=1
)

print("\nTop cluster pairs (raw LR):")
display(
    lr_df["cluster_pair"]
    .value_counts()
    .head(10)
    .rename_axis("cluster_pair")
    .reset_index(name="n_lr")
)

# -----------------------------------
# Sanity flag
# -----------------------------------
print("\n=== QC STATUS: OK ===")

# %% [markdown]
## Cohort-level aggregation

Aggregate per-sample outputs into combined tables for the full cohort.

# %% [code]
# Combine all BREAST sample summaries

import pandas as pd
from pathlib import Path
from IPython.display import display

BASE_DIR = Path("/Users/sally/Desktop/Metrics_project/Samples/GSE243022_RAW")

samples = [
    "GSM7777520_A1",
    "GSM7777521_B1",
    "GSM7777522_C1",
    "GSM7777523_D1",
]

dfs = []

for s in samples:
    f = BASE_DIR / s / "boundary_enriched_outputs" / f"{s}_sample_summary.csv"
    if not f.exists():
        raise FileNotFoundError(f"Missing summary: {f}")
    df = pd.read_csv(f)
    dfs.append(df)

combined = pd.concat(dfs, ignore_index=True)

print("=== BREAST SAMPLE SUMMARIES ===")
display(combined)

print("\n=== REGIME COUNTS ===")
display(
    combined["regime"]
    .value_counts()
    .rename_axis("regime")
    .reset_index(name="n")
)

print("\n=== CONSISTENCY CHECK ===")
unique_regimes = combined["regime"].unique()
print("Unique regimes:", unique_regimes)
print("Consistent:", len(unique_regimes) == 1)

# %% [markdown]
## Export verification

Display final export paths and verify that supplementary outputs were written successfully.

# %% [code]
print("=== BREAST REPLICATION EXPORT ===")

lr_row_path = OUT_DIR / f"{SAMPLE_NAME}_lr_row_table.csv"
lr_collapsed_path = OUT_DIR / f"{SAMPLE_NAME}_lr_collapsed_table.csv"
sample_summary_path = OUT_DIR / f"{SAMPLE_NAME}_sample_summary.csv"

lr_full.to_csv(lr_row_path, index=False)
lr_collapsed.to_csv(lr_collapsed_path, index=False)
sample_summary.to_csv(sample_summary_path, index=False)

print("Saved:")
print(" -", lr_row_path.name)
print(" -", lr_collapsed_path.name)
print(" -", sample_summary_path.name)
```
