## Supplementary Notebook 1-5 for "Geometry-aware ligand-receptor analysis distinguishes interface association from spatial localization and reveals a continuum of tumor communication": Figures.ipynb

```
# %% [markdown]
# Publication-ready figures for the spatial ligand–receptor manuscript

This notebook is a polished, reproducible figure-production notebook built from the merged manuscript tables:

- `all_lr_rows.csv`
- `all_lr_collapsed.csv`
- `all_sample_summaries.csv`

It centralizes configuration, styling, helper functions, QC, and figure export to PNG and PDF.


# %% [code]

from pathlib import Path
import pandas as pd
import numpy as np
import matplotlib.pyplot as plt
from matplotlib.patches import Patch
from pandas.errors import EmptyDataError

# =========================
# Configuration
# =========================
PROJECT_DIR = Path("/Users/sally/Desktop/Metrics_project")
TABLE_DIR = PROJECT_DIR / "FINAL_MANUSCRIPT_TABLES"
FIG_DIR = PROJECT_DIR / "FINAL_MANUSCRIPT_FIGURES_PUBLICATION"
MANIFEST_DIR = PROJECT_DIR / "FINAL_MANUSCRIPT_FIGURE_MANIFEST"
FIG_DIR.mkdir(exist_ok=True)
MANIFEST_DIR.mkdir(exist_ok=True)

# =========================
# Publication-style defaults
# =========================
plt.rcParams.update({
    "figure.dpi": 120,
    "savefig.dpi": 300,
    "font.size": 10,
    "axes.titlesize": 12,
    "axes.labelsize": 10,
    "axes.linewidth": 0.8,
    "xtick.labelsize": 9,
    "ytick.labelsize": 9,
    "legend.fontsize": 9,
    "legend.title_fontsize": 9,
    "pdf.fonttype": 42,   # editable text in Illustrator
    "ps.fonttype": 42,
})

TISSUE_ORDER = ["breast", "colorectal", "melanoma", "pdac"]
TISSUE_COLORS = {
    "breast": "#4477AA",
    "colorectal": "#EE7733",
    "melanoma": "#228833",
    "pdac": "#CC3311",
    "unknown": "#999999",
}
REGIME_ORDER = ["localized", "mixed", "diffuse"]
REGIME_COLORS = {
    "localized": "#66CCEE",
    "mixed": "#AA3377",
    "diffuse": "#BBBBBB",
}

def fix_tissue(sample_id):
    s = str(sample_id).lower()
    if "pdac" in s:
        return "pdac"
    if "gsm798" in s or "melanoma" in s:
        return "melanoma"
    if "gsm671" in s or "crc" in s:
        return "colorectal"
    if "gsm777" in s or "section" in s:
        return "breast"
    return "unknown"

def add_tissue(df):
    out = df.copy()
    if "sample_id" not in out.columns:
        raise KeyError("Expected a sample_id column.")
    out["tissue"] = out["sample_id"].apply(fix_tissue)
    return out

def save_figure(fig, stem):
    png_path = FIG_DIR / f"{stem}.png"
    pdf_path = FIG_DIR / f"{stem}.pdf"
    fig.savefig(png_path, bbox_inches="tight")
    fig.savefig(pdf_path, bbox_inches="tight")
    print("Saved:")
    print(" -", png_path)
    print(" -", pdf_path)
    return png_path, pdf_path

def load_csv(path):
    try:
        return pd.read_csv(path)
    except EmptyDataError:
        raise RuntimeError(f"CSV is empty: {path}")

def normalize_regime_columns(df):
    out = df.copy()
    for col in REGIME_ORDER:
        if col not in out.columns:
            out[col] = 0.0
    return out[REGIME_ORDER]


# %% [code]

# =========================
# Load merged manuscript tables
# =========================
all_lr_rows = add_tissue(load_csv(TABLE_DIR / "all_lr_rows.csv"))
all_lr_collapsed = add_tissue(load_csv(TABLE_DIR / "all_lr_collapsed.csv"))
all_sample_summaries = add_tissue(load_csv(TABLE_DIR / "all_sample_summaries.csv"))

print("Loaded:")
print(" - all_lr_rows:", all_lr_rows.shape)
print(" - all_lr_collapsed:", all_lr_collapsed.shape)
print(" - all_sample_summaries:", all_sample_summaries.shape)


# %% [code]

# =========================
# Quick QC
# =========================
print("Row table tissues:")
print(all_lr_rows["tissue"].value_counts(dropna=False).sort_index(), end="\n\n")

print("Collapsed table tissues:")
print(all_lr_collapsed["tissue"].value_counts(dropna=False).sort_index(), end="\n\n")

print("Sample summary tissues:")
print(all_sample_summaries["tissue"].value_counts(dropna=False).sort_index(), end="\n\n")

print("Sample-level regimes:")
if "regime" in all_sample_summaries.columns:
    print(all_sample_summaries["regime"].value_counts(dropna=False))
else:
    print("No regime column found.")


# %% [markdown]
## Figure 2A — Top pathways per tissue by mean boundary score

# %% [code]

df = all_lr_rows.copy()
df = df.dropna(subset=["pathway_name", "boundary_score", "sample_id"])
df = df[df["pathway_name"].astype(str).str.strip() != ""]
df = df[df["tissue"] != "unknown"]

pathway_summary = (
    df.groupby(["tissue", "pathway_name"], as_index=False)
      .agg(
          mean_boundary=("boundary_score", "mean"),
          n_lr_rows=("interaction_name", "size"),
          n_samples=("sample_id", "nunique"),
      )
      .sort_values(
          ["tissue", "mean_boundary", "n_samples", "n_lr_rows"],
          ascending=[True, False, False, False]
      )
)

top_n = 10
top_pathways = pathway_summary.groupby("tissue", group_keys=False).head(top_n).copy()

fig, axes = plt.subplots(len(TISSUE_ORDER), 1, figsize=(10, 12), squeeze=False)

for ax, tissue in zip(axes.flatten(), TISSUE_ORDER):
    sub = top_pathways[top_pathways["tissue"] == tissue].sort_values("mean_boundary", ascending=True)
    ax.barh(sub["pathway_name"], sub["mean_boundary"], color=TISSUE_COLORS[tissue], alpha=0.9)
    ax.set_title(f"{tissue.capitalize()}")
    ax.set_xlabel("Mean boundary score")
    ax.set_ylabel("Pathway")

fig.suptitle("Figure 2A. Top pathways per tissue", y=1.01)
plt.tight_layout()
save_figure(fig, "Figure_2A_top_pathways_by_tissue_publication")
plt.show()


# %% [markdown]
## Figure 2B — Regime composition of recurrent pathways

# %% [code]

df = all_lr_rows.copy()
df = df.dropna(subset=["pathway_name", "interaction_regime"])
df = df[df["pathway_name"].astype(str).str.strip() != ""]

top_paths = (
    df.groupby("pathway_name")
      .size()
      .sort_values(ascending=False)
      .head(15)
      .index
)

plot_df = df[df["pathway_name"].isin(top_paths)].copy()

regime_counts = (
    plot_df.groupby(["pathway_name", "interaction_regime"])
           .size()
           .unstack(fill_value=0)
)

regime_frac = normalize_regime_columns(regime_counts).div(regime_counts.sum(axis=1), axis=0)
regime_frac = regime_frac.sort_values(by="diffuse", ascending=False)

fig, ax = plt.subplots(figsize=(12, 5))
bottom = np.zeros(len(regime_frac))
x = np.arange(len(regime_frac))

for regime in REGIME_ORDER:
    vals = regime_frac[regime].values
    ax.bar(x, vals, bottom=bottom, label=regime, color=REGIME_COLORS[regime], width=0.85)
    bottom += vals

ax.set_xticks(x)
ax.set_xticklabels(regime_frac.index, rotation=45, ha="right")
ax.set_ylabel("Fraction of LR rows")
ax.set_xlabel("Pathway")
ax.set_title("Figure 2B. Regime composition of recurrent pathways")
ax.legend(title="Interaction regime", bbox_to_anchor=(1.02, 1), loc="upper left")

plt.tight_layout()
save_figure(fig, "Figure_2B_regime_composition_recurrent_pathways_publication")
plt.show()


# %% [markdown]
## Figure 3A — Observed boundary score vs null mean

# %% [code]

df = all_lr_collapsed.copy()
df = df.dropna(subset=["boundary_score", "null_mean"])
df = df[df["tissue"] != "unknown"]

fig, ax = plt.subplots(figsize=(6.2, 6.2))

for tissue in TISSUE_ORDER:
    sub = df[df["tissue"] == tissue]
    ax.scatter(
        sub["null_mean"],
        sub["boundary_score"],
        s=8,
        alpha=0.25,
        label=tissue,
        color=TISSUE_COLORS[tissue]
    )

lims = [
    min(df["null_mean"].min(), df["boundary_score"].min()),
    max(df["null_mean"].max(), df["boundary_score"].max()),
]
ax.plot(lims, lims, linestyle="--", linewidth=1.0, color="#444444")
ax.set_xlabel("Null mean")
ax.set_ylabel("Observed boundary score")
ax.set_title("Figure 3A. Observed vs null interface enrichment")
ax.legend(title="Tissue", frameon=False)

plt.tight_layout()
save_figure(fig, "Figure_3A_observed_vs_null_publication")
plt.show()


# %% [markdown]
## Figure 3B — Null gap distribution

# %% [code]

df = all_lr_collapsed.copy()
df = df.dropna(subset=["null_gap"])
df = df[df["tissue"] != "unknown"]

fig, ax = plt.subplots(figsize=(7, 5))

for tissue in TISSUE_ORDER:
    sub = df[df["tissue"] == tissue]
    ax.hist(
        sub["null_gap"],
        bins=60,
        density=True,
        histtype="step",
        linewidth=1.8,
        label=tissue,
        color=TISSUE_COLORS[tissue]
    )

ax.axvline(0, linestyle="--", linewidth=1.0, color="#444444")
ax.set_xlabel("Null gap (boundary_score - null_mean)")
ax.set_ylabel("Density")
ax.set_title("Figure 3B. Distribution of null gaps")
ax.legend(title="Tissue", frameon=False)

plt.tight_layout()
save_figure(fig, "Figure_3B_null_gap_overlay_publication")
plt.show()


# %% [markdown]
## Figure 3C — Effect size vs significance

# %% [code]

df = all_lr_collapsed.copy()
df = df.dropna(subset=["boundary_score", "empirical_p"])
df = df[df["tissue"] != "unknown"].copy()
df["empirical_p"] = df["empirical_p"].clip(lower=1e-6)

fig, ax = plt.subplots(figsize=(6.2, 6.2))

for tissue in TISSUE_ORDER:
    sub = df[df["tissue"] == tissue]
    ax.scatter(
        sub["boundary_score"],
        -np.log10(sub["empirical_p"]),
        s=8,
        alpha=0.25,
        label=tissue,
        color=TISSUE_COLORS[tissue]
    )

ax.set_xlabel("Boundary score")
ax.set_ylabel("-log10(empirical p-value)")
ax.set_title("Figure 3C. Effect size vs significance")
ax.legend(title="Tissue", frameon=False)

plt.tight_layout()
save_figure(fig, "Figure_3C_effect_vs_significance_publication")
plt.show()


# %% [markdown]
## Figure 4A — Boundary score vs geometry-aware score

# %% [code]

df = all_lr_collapsed.copy()
df = df.dropna(subset=["boundary_score", "geometry_score"])
df = df[df["tissue"] != "unknown"]

fig, ax = plt.subplots(figsize=(6.2, 6.2))

for tissue in TISSUE_ORDER:
    sub = df[df["tissue"] == tissue]
    ax.scatter(
        sub["boundary_score"],
        sub["geometry_score"],
        s=8,
        alpha=0.25,
        label=tissue,
        color=TISSUE_COLORS[tissue]
    )

lims = [
    min(df["boundary_score"].min(), df["geometry_score"].min()),
    max(df["boundary_score"].max(), df["geometry_score"].max()),
]
ax.plot(lims, lims, linestyle="--", linewidth=1.0, color="#444444")
ax.set_xlabel("Boundary score")
ax.set_ylabel("Geometry-aware score")
ax.set_title("Figure 4A. Boundary versus geometry-aware scores")
ax.legend(title="Tissue", frameon=False)

plt.tight_layout()
save_figure(fig, "Figure_4A_boundary_vs_geometry_publication")
plt.show()


# %% [markdown]
## Figure 4B — Retention distribution

# %% [code]

df = all_lr_collapsed.copy()
df = df.dropna(subset=["retention"])
df = df[df["tissue"] != "unknown"]
df = df[df["retention"] >= 0]

fig, ax = plt.subplots(figsize=(7, 5))

for tissue in TISSUE_ORDER:
    sub = df[df["tissue"] == tissue]
    ax.hist(
        sub["retention"],
        bins=60,
        density=True,
        histtype="step",
        linewidth=1.8,
        label=tissue,
        color=TISSUE_COLORS[tissue]
    )

ax.set_xlabel("Retention (geometry_score / boundary_score)")
ax.set_ylabel("Density")
ax.set_title("Figure 4B. Retention distribution")
ax.legend(title="Tissue", frameon=False)

plt.tight_layout()
save_figure(fig, "Figure_4B_retention_distribution_publication")
plt.show()


# %% [markdown]
## Figure 4C — Sample-level retention

# %% [code]

df = all_sample_summaries.copy()
df = df.dropna(subset=["retention", "sample_id"]).copy()
df = df[df["tissue"] != "unknown"].copy()
df = df.sort_values("retention", ascending=False).reset_index(drop=True)

fig, ax = plt.subplots(figsize=(11, 5.5))
x = np.arange(len(df))
bar_colors = [TISSUE_COLORS.get(t, TISSUE_COLORS["unknown"]) for t in df["tissue"]]

ax.bar(x, df["retention"], color=bar_colors, width=0.8)
ax.set_xticks(x)
ax.set_xticklabels(df["sample_id"], rotation=90)
ax.set_xlabel("Sample")
ax.set_ylabel("Retention")
ax.set_title("Figure 4C. Sample-level retention")

legend_handles = [Patch(facecolor=TISSUE_COLORS[t], label=t) for t in TISSUE_ORDER]
ax.legend(handles=legend_handles, title="Tissue", frameon=False)

plt.tight_layout()
save_figure(fig, "Figure_4C_sample_level_retention_publication")
plt.show()


# %% [markdown]
## Figure 5A — Retention vs fraction diffuse

# %% [code]

df = all_sample_summaries.copy()
df = df.dropna(subset=["retention", "frac_diffuse", "sample_id"]).copy()
df = df[df["tissue"] != "unknown"].copy()

fig, ax = plt.subplots(figsize=(7, 6))

for tissue in TISSUE_ORDER:
    sub = df[df["tissue"] == tissue]
    ax.scatter(
        sub["retention"],
        sub["frac_diffuse"],
        s=75,
        alpha=0.9,
        color=TISSUE_COLORS[tissue],
        label=tissue
    )

ax.set_xlabel("Retention")
ax.set_ylabel("Fraction diffuse")
ax.set_title("Figure 5A. Retention versus diffuse fraction")
ax.legend(title="Tissue", frameon=False)

plt.tight_layout()
save_figure(fig, "Figure_5A_retention_vs_fraction_diffuse_publication")
plt.show()


# %% [markdown]
## Figure 5B — Retention vs null gap fraction

# %% [code]

df = all_sample_summaries.copy()
df = df.dropna(subset=["retention", "null_gap_fraction"]).copy()
df = df[df["tissue"] != "unknown"].copy()

fig, ax = plt.subplots(figsize=(7, 6))

for tissue in TISSUE_ORDER:
    sub = df[df["tissue"] == tissue]
    ax.scatter(
        sub["retention"],
        sub["null_gap_fraction"],
        s=75,
        alpha=0.9,
        color=TISSUE_COLORS[tissue],
        label=tissue
    )

ax.set_xlabel("Retention")
ax.set_ylabel("Null gap fraction")
ax.set_title("Figure 5B. Retention versus null-related metric")
ax.legend(title="Tissue", frameon=False)

plt.tight_layout()
save_figure(fig, "Figure_5B_retention_vs_null_gap_fraction_publication")
plt.show()


# %% [markdown]
## Figure 5C — Retention by tissue

# %% [code]

df = all_sample_summaries.copy()
df = df.dropna(subset=["retention"]).copy()
df = df[df["tissue"] != "unknown"].copy()

data = [df.loc[df["tissue"] == t, "retention"].values for t in TISSUE_ORDER]

fig, ax = plt.subplots(figsize=(7, 5))
bp = ax.boxplot(data, labels=TISSUE_ORDER, patch_artist=True)

for patch, tissue in zip(bp["boxes"], TISSUE_ORDER):
    patch.set_facecolor(TISSUE_COLORS[tissue])
    patch.set_alpha(0.6)

ax.set_xlabel("Tissue")
ax.set_ylabel("Retention")
ax.set_title("Figure 5C. Retention by tissue")

plt.tight_layout()
save_figure(fig, "Figure_5C_retention_by_tissue_publication")
plt.show()


# %% [markdown]
## Figure 5D — Sample-level regime outcome

# %% [code]

df = all_sample_summaries.copy()
df = df.dropna(subset=["sample_id", "regime"]).copy()
df = df[df["tissue"] != "unknown"].copy()

tissue_rank = {t: i for i, t in enumerate(TISSUE_ORDER)}
df["tissue_rank"] = df["tissue"].map(tissue_rank)
df = df.sort_values(["tissue_rank", "sample_id"]).reset_index(drop=True)

regime_to_num = {r: i for i, r in enumerate(REGIME_ORDER)}
df["regime_num"] = df["regime"].map(regime_to_num)

fig, ax = plt.subplots(figsize=(11, 4))
x = np.arange(len(df))
colors = [TISSUE_COLORS.get(t, TISSUE_COLORS["unknown"]) for t in df["tissue"]]

ax.scatter(x, df["regime_num"], c=colors, s=80)
ax.set_xticks(x)
ax.set_xticklabels(df["sample_id"], rotation=90)
ax.set_yticks(range(len(REGIME_ORDER)))
ax.set_yticklabels(REGIME_ORDER)
ax.set_xlabel("Sample")
ax.set_ylabel("Sample-level regime")
ax.set_title("Figure 5D. Sample-level regime outcome")

legend_handles = [Patch(facecolor=TISSUE_COLORS[t], label=t) for t in TISSUE_ORDER]
ax.legend(handles=legend_handles, title="Tissue", bbox_to_anchor=(1.02, 1), loc="upper left", frameon=False)

plt.tight_layout()
save_figure(fig, "Figure_5D_sample_level_regime_outcome_publication")
plt.show()


# %% [markdown]
## Figure 6A — Mean retention of conserved pathways across tissues

# %% [code]

df = all_lr_rows.copy()
df = df.dropna(subset=["pathway_name", "retention", "interaction_name"]).copy()
df = df[df["tissue"] != "unknown"].copy()
df = df[df["pathway_name"].astype(str).str.strip() != ""]

pathway_tissue = (
    df.groupby(["pathway_name", "tissue"], as_index=False)
      .agg(
          mean_retention=("retention", "mean"),
          n_rows=("interaction_name", "size"),
          n_samples=("sample_id", "nunique")
      )
)

pathway_recurrence = (
    pathway_tissue.groupby("pathway_name", as_index=False)
                 .agg(
                     n_tissues=("tissue", "nunique"),
                     total_rows=("n_rows", "sum"),
                     total_samples=("n_samples", "sum")
                 )
)

top_paths = (
    pathway_recurrence
    .sort_values(["n_tissues", "total_samples", "total_rows"], ascending=[False, False, False])
    .head(15)["pathway_name"]
    .tolist()
)

plot_df = pathway_tissue[pathway_tissue["pathway_name"].isin(top_paths)].copy()
heatmap_df = plot_df.pivot(index="pathway_name", columns="tissue", values="mean_retention").reindex(columns=TISSUE_ORDER)
heatmap_df = heatmap_df.loc[heatmap_df.mean(axis=1).sort_values(ascending=False).index]

fig, ax = plt.subplots(figsize=(7, 8))
im = ax.imshow(heatmap_df.values, aspect="auto", interpolation="nearest")
ax.set_xticks(np.arange(len(heatmap_df.columns)))
ax.set_xticklabels(heatmap_df.columns)
ax.set_yticks(np.arange(len(heatmap_df.index)))
ax.set_yticklabels(heatmap_df.index)
ax.set_xlabel("Tissue")
ax.set_ylabel("Pathway")
ax.set_title("Figure 6A. Pathway retention across tissues")

cbar = fig.colorbar(im, ax=ax)
cbar.set_label("Mean retention")

plt.tight_layout()
save_figure(fig, "Figure_6A_conserved_pathway_retention_heatmap_publication")
plt.show()


# %% [markdown]
## Figure 6B — Regime composition of conserved pathways

# %% [code]

df = all_lr_rows.copy()
df = df.dropna(subset=["pathway_name", "interaction_regime"]).copy()
df = df[df["tissue"] != "unknown"].copy()
df = df[df["pathway_name"].astype(str).str.strip() != ""]

pathway_tissue = (
    df.groupby(["pathway_name", "tissue"], as_index=False)
      .agg(
          n_rows=("interaction_name", "size"),
          n_samples=("sample_id", "nunique")
      )
)

pathway_recurrence = (
    pathway_tissue.groupby("pathway_name", as_index=False)
                 .agg(
                     n_tissues=("tissue", "nunique"),
                     total_rows=("n_rows", "sum"),
                     total_samples=("n_samples", "sum")
                 )
)

top_paths = (
    pathway_recurrence
    .sort_values(["n_tissues", "total_samples", "total_rows"], ascending=[False, False, False])
    .head(15)["pathway_name"]
    .tolist()
)

plot_df = df[df["pathway_name"].isin(top_paths)].copy()
regime_counts = (
    plot_df.groupby(["pathway_name", "interaction_regime"])
           .size()
           .unstack(fill_value=0)
)
regime_frac = normalize_regime_columns(regime_counts).div(regime_counts.sum(axis=1), axis=0).sort_values(by="diffuse", ascending=False)

fig, ax = plt.subplots(figsize=(12, 5))
bottom = np.zeros(len(regime_frac))
x = np.arange(len(regime_frac))

for regime in REGIME_ORDER:
    vals = regime_frac[regime].values
    ax.bar(x, vals, bottom=bottom, width=0.85, label=regime, color=REGIME_COLORS[regime])
    bottom += vals

ax.set_xticks(x)
ax.set_xticklabels(regime_frac.index, rotation=45, ha="right")
ax.set_xlabel("Pathway")
ax.set_ylabel("Fraction of LR rows")
ax.set_title("Figure 6B. Regime composition of conserved pathways")
ax.legend(title="Interaction regime", bbox_to_anchor=(1.02, 1), loc="upper left", frameon=False)

plt.tight_layout()
save_figure(fig, "Figure_6B_conserved_pathway_regime_composition_publication")
plt.show()


# %% [markdown]
## Figure 6C — Tissue-specific retention of conserved pathways

# %% [code]

df = all_lr_rows.copy()
df = df.dropna(subset=["pathway_name", "tissue", "retention", "interaction_name"]).copy()
df = df[df["tissue"] != "unknown"].copy()
df = df[df["pathway_name"].astype(str).str.strip() != ""]

pathway_tissue = (
    df.groupby(["pathway_name", "tissue"], as_index=False)
      .agg(
          n_rows=("interaction_name", "size"),
          n_samples=("sample_id", "nunique")
      )
)

pathway_recurrence = (
    pathway_tissue.groupby("pathway_name", as_index=False)
                 .agg(
                     n_tissues=("tissue", "nunique"),
                     total_rows=("n_rows", "sum"),
                     total_samples=("n_samples", "sum")
                 )
)

top_paths = (
    pathway_recurrence
    .sort_values(["n_tissues", "total_samples", "total_rows"], ascending=[False, False, False])
    .head(12)["pathway_name"]
    .tolist()
)

summary_df = (
    df[df["pathway_name"].isin(top_paths)]
      .groupby(["pathway_name", "tissue"], as_index=False)
      .agg(mean_retention=("retention", "mean"))
)

fig, ax = plt.subplots(figsize=(14, 6))
x = np.arange(len(top_paths))
width = 0.2

for i, tissue in enumerate(TISSUE_ORDER):
    sub = summary_df[summary_df["tissue"] == tissue].set_index("pathway_name").reindex(top_paths).reset_index()
    xpos = x + (i - 1.5) * width
    ax.bar(xpos, sub["mean_retention"], width=width, label=tissue, color=TISSUE_COLORS[tissue])

ax.set_xticks(x)
ax.set_xticklabels(top_paths, rotation=45, ha="right")
ax.set_xlabel("Pathway")
ax.set_ylabel("Mean retention")
ax.set_title("Figure 6C. Tissue-level comparison of pathway retention")
ax.legend(title="Tissue", frameon=False)

plt.tight_layout()
save_figure(fig, "Figure_6C_tissue_specific_pathway_retention_publication")
plt.show()


# %% [markdown]
## Figure 6D — Global regime composition across tissues

# %% [code]

df = all_lr_rows.copy()
df = df.dropna(subset=["interaction_regime", "tissue"]).copy()
df = df[df["tissue"] != "unknown"].copy()

regime_counts = (
    df.groupby(["tissue", "interaction_regime"])
      .size()
      .unstack(fill_value=0)
      .reindex(index=TISSUE_ORDER)
)
regime_frac = normalize_regime_columns(regime_counts).div(regime_counts.sum(axis=1), axis=0)

fig, ax = plt.subplots(figsize=(7, 5))
bottom = np.zeros(len(regime_frac))
x = np.arange(len(regime_frac))

for regime in REGIME_ORDER:
    vals = regime_frac[regime].values
    ax.bar(x, vals, bottom=bottom, width=0.8, label=regime, color=REGIME_COLORS[regime])
    bottom += vals

ax.set_xticks(x)
ax.set_xticklabels(regime_frac.index)
ax.set_xlabel("Tissue")
ax.set_ylabel("Fraction of LR rows")
ax.set_title("Figure 6D. Global regime composition by tissue")
ax.legend(title="Interaction regime", bbox_to_anchor=(1.02, 1), loc="upper left", frameon=False)

plt.tight_layout()
save_figure(fig, "Figure_6D_global_regime_by_tissue_publication")
plt.show()


# %% [markdown]
## Figure and table manifest

# %% [code]

figure_files = []
for ext in ("*.png", "*.pdf"):
    for f in sorted(FIG_DIR.glob(ext)):
        figure_files.append({
            "filename": f.name,
            "format": f.suffix.replace(".", ""),
            "path": str(f),
            "size_bytes": f.stat().st_size
        })

table_files = []
for f in sorted(TABLE_DIR.glob("*.csv")):
    table_files.append({
        "filename": f.name,
        "path": str(f),
        "size_bytes": f.stat().st_size
    })

figure_manifest = pd.DataFrame(figure_files).sort_values(["filename", "format"])
table_manifest = pd.DataFrame(table_files).sort_values("filename")

figure_manifest.to_csv(MANIFEST_DIR / "figure_manifest.csv", index=False)
table_manifest.to_csv(MANIFEST_DIR / "table_manifest.csv", index=False)

print("Saved:")
print(" -", MANIFEST_DIR / "figure_manifest.csv")
print(" -", MANIFEST_DIR / "table_manifest.csv")

figure_manifest.head()
```
